# Identification of trans-AT polyketide clusters in two marine bacteria reveals cryptic similarities between distinct symbiosis factors

**DOI:** 10.1101/2020.09.18.303172

**Authors:** Dina Kačar, Librada M Cañedo, Pilar Rodríguez, Elena Gonzalez, Beatriz Galán, Carmen Schleissner, Stefan Leopold-Messer, Jörn Piel, Carmen Cuevas, Fernando de la Calle, José L. García

## Abstract

Glutaramide-containing polyketides are known as potent antitumoral and antimetastatic agents. However, the associated gene clusters have only been identified and studied in a few *Streptomyces* producers and sole *Burkholderia gladioli* symbiont. The new glutaramide-family polyketides, denominated sesbanimides D, E and F along with the previously known sesbanimide A and C, were isolated from two marine alphaproteobacteria *Stappia indica* PHM037 and *Labrenzia aggregata* PHM038. Structures of the isolated compounds were elucidated based on 1D and 2D homo and heteronuclear NMR analyses and ESI-MS spectrometry. All compounds exhibited strong antitumor activity in lung, breast and colorectal cancer cell lines. Subsequent whole genome sequencing and genome mining revealed the presence of the *trans*-AT PKS gene cluster responsible for the sesbanimide biosynthesis, described as *sbn* cluster, and the sesbanimide modular assembly is proposed. Interestingly, numerous homologous orphan gene clusters were localized in distantly related bacteria and used as comparative genomic assets for a more global characterization of *sbn* like-clusters. Strikingly, the modular architecture of downstream mixed type PKS/NRPS, SbnQ, revealed high similarity to PedH in pederin and Lab13 in labrenzin gene clusters, although those clusters are responsible for the production of structurally completely different molecules. The unexpected presence of SbnQ homologs in unrelated polyketide gene clusters across phylogenetically distant bacteria, raises intriguing questions about the evolutionary relationship between glutaramide-like and pederin-like pathways, as well as the functionality of their synthetic products.

**Significance:** Glutaramide-containing polyketides are still a largely understudied group of polyketides, produced mainly by the genera *Streptomyces*, with a great potential for antitumor drug production. Here, we describe genomes of two cultivable marine bacteria, *Stappia indica* PHM037 and *Labrenzia aggregata* PHM038, producers of the cytotoxic glutaramide-family polyketides sesbanimide A and C with chemical elucidation of newly identified analogs D, E and F. Genome mining revealed *trans*-AT PKS gene cluster responsible for sesbanimide biosynthesis. Although there are numerous homologous gene clusters present in remarkably different bacteria, this is the first time that the biosynthesis product has been reported. The comparative genome analysis reveals stunning, cryptic evolutionary relationship between sesbanimides, glutaramides from *Streptomyces* spp. and the pederin-family gene clusters.

---

Seeds of *Sesbania* species, commonly referred to as rattlebush, rattlebox, or coffeebean, have been known to be toxic to livestock for centuries (1). Extractions of *Sesbania drummondii* (Rydb.) seeds, a shrub inhabiting the coastal plains between Florida and Texas, revealed sesbanimides as the toxic compounds (2). The first cytotoxic activity of the extract *in vivo* in tumor cell lines was shown for ethanolic extracts of *Sesbania* seeds (3, 4). Initially, the activity was attributed to the previously identified compounds sesbanine and drummondol, although the antitumor activity was absent after HPLC purification steps (3). Further fractionations of the previously reported extracts and separation from the sesbanine, elucidated the tricyclic structure of sesbanimide A (2). Later on, full details of chemical identification of sesbanimide A, B and C were revealed (5). Nonetheless, little is known about their origin and biosynthesis.

The molecular structure of sesbanimides is based on the glutaramide moiety, which is common in glutaramide-containig polyketides well-studied for their antitumor properties. Examples of related polyketides are *iso*-migrastatin, migrastatin, dorrigocins, lactimidomycin and analogs reported as potent antimetastatic agents due to their cell migration inhibition activity (6, 7). Some of these analogs also exhibit protein synthesis inhibitory activity by blocking the translation elongation, suggesting the glutarimide polyketides play an essential role in ribosome binding (8). Other examples of structurally related glutarimide-containing polyketides are cycloheximide that has been used for decades as inhibitor of eukaryotic protein synthesis (9, 10) and 9-methylstreptimidone with potential antiviral activity by exerting a protective effect against influenza A2 (H2N2) virus in mice (11). Recently, two new methylstreptimidone analogs have been isolated showing moderate antitumor activity (12). Apart from antitumor activity against human glioma, streptoglutarimides isolated from a marine actinomycete have also shown antibacterial and antifungal properties (13).

Known glutarimide polyketides have been isolated from actinomycetes (12–16) with the exception of recently identified gladiofungin from *Burkholderia gladioli* (17). Nevertheless, the production of the sesbanimide analogs has been attributed to Gram-negative bacteria, *i.e*., marine *Agrobacterium* strains, PH-103 and PH-A034C, isolated from the tunicates *Ecteinascidia turbinata* collected in Florida and *Polycitonidae* sp. collected in Turkey, that produced sesbanimide A and C, respectively (18). Until recently, no other producers have been reported and the genetic and metabolic context behind the sesbanimide biosynthesis has remained unknown.

Herein, we describe the isolation of new analogues of sesbanimides produced by two alphaproteobacteria and the identification of their biosynthetic gene clusters. Genome sequencing and annotation of these strains have allowed us to identify the *trans*-acyltransferase polyketide synthase (*trans*-AT PKS) gene clusters associated with sesbanimide production and to propose the biosynthesis pathway according to the ascribed gene/enzyme functionalities resulting from bioinformatics and comparative studies. The genomic analyses reveal surprising insights into the biosynthesis of these polyketides as well as the wide distribution of the biosynthetic genes among phylogenetically distant microorganisms and diverse ecological niches.

## Results and Discussion

### Bacterial cultivation and identification of sesbanimide analogs

Cultures of the isolated strains *Stappia indica* PHM037 and *Labrenzia aggregata* PHM038 were scaled-up and their crude extracts were fractionated by preparative HPLC to isolate the active compounds **1** to **5** (Fig. 1). The structures of these compounds were elucidated by a combination of spectroscopic methods. Sesbanimide A (**1**) and its newly identified acyclic form sesbanimide D (**2**) were produced by strain PHM038; and sesbanimide C (**3**) and its new acyclic analog sesbanimide E (**4**) together with a novel sesbanimide F (**5**) were produced by the strain PHM037 (Fig. 1). Compounds **1** and **3** were identified by NMR and mass spectra as sesbanimides A and C previously isolated from two marine strains PH-103 and PH-A034C, respectively, closely related to *Agrobacterium* species (18). Compounds **2** (ESIMS *m/z* 350.1 [M+Na]^+^ and 677.3 [2M+Na]^+^; HRESIMS *m/z* 350.1244 [M+Na]^+^ (calcd. for C_15_H_21_NO_7_Na, 350.1220) and **4** (ESIMS *m/z* 336.1 [M+Na]^+^ and 649.3 [2M+Na]^+^; HRESIMS *m/z* 336.1400 [M+Na]^+^ (calcd. for C_15_H_23_NO_6_Na, 336.1418) possess the same molecular weight and molecular formula as compounds **1** and **3**, respectively. The structures of the new compounds **2** and **4** were elucidated using 1D and 2D homo- and hetero-nuclear NMR analyses and both compounds showed a ketone carbonyl at C10 instead of the hemiketal in **1** and **3** with a furoketal ring structure, the rest of NMR data were very similar to those described for sesbanimides A and C (18). These data suggested that in compounds **2** and **4** the furoketal ring was opened and were identified as acyclic keto analogs at C10 of **1** and **3** respectively. The HMBC spectra of **2** and **4** demonstrated the expected key correlations. Compound **5** has a molecular formula C_26_H_37_NO_9_ deduced from its ESIMS *m/z* 508.3 [M+H]^+^ and 530.3 [M+Na]^+^; HRESIMS *m/z* 530.2363 [M+Na]^+^ (calcd. for C_26_H_37_NO_9_Na, 530.2361) as well as its ^13^C NMR data. Detailed 2D NMR spectroscopic analyses indicated that **5** possessed the same core structure as **4** linked at C13 with a dicarboxylic unsaturated eleven-carbon chain through esterification. The HMBC spectra of **5** demonstrated the expected key correlations. All related NMR and ESIMS spectra are shown in Figs. S1-S24 in supporting information. According to the structural relationships of compounds **2**, **4** and **5** with sesbanimides A and C, they are referred as sesbanimides D, E and F, respectively.

**Figure 1.**
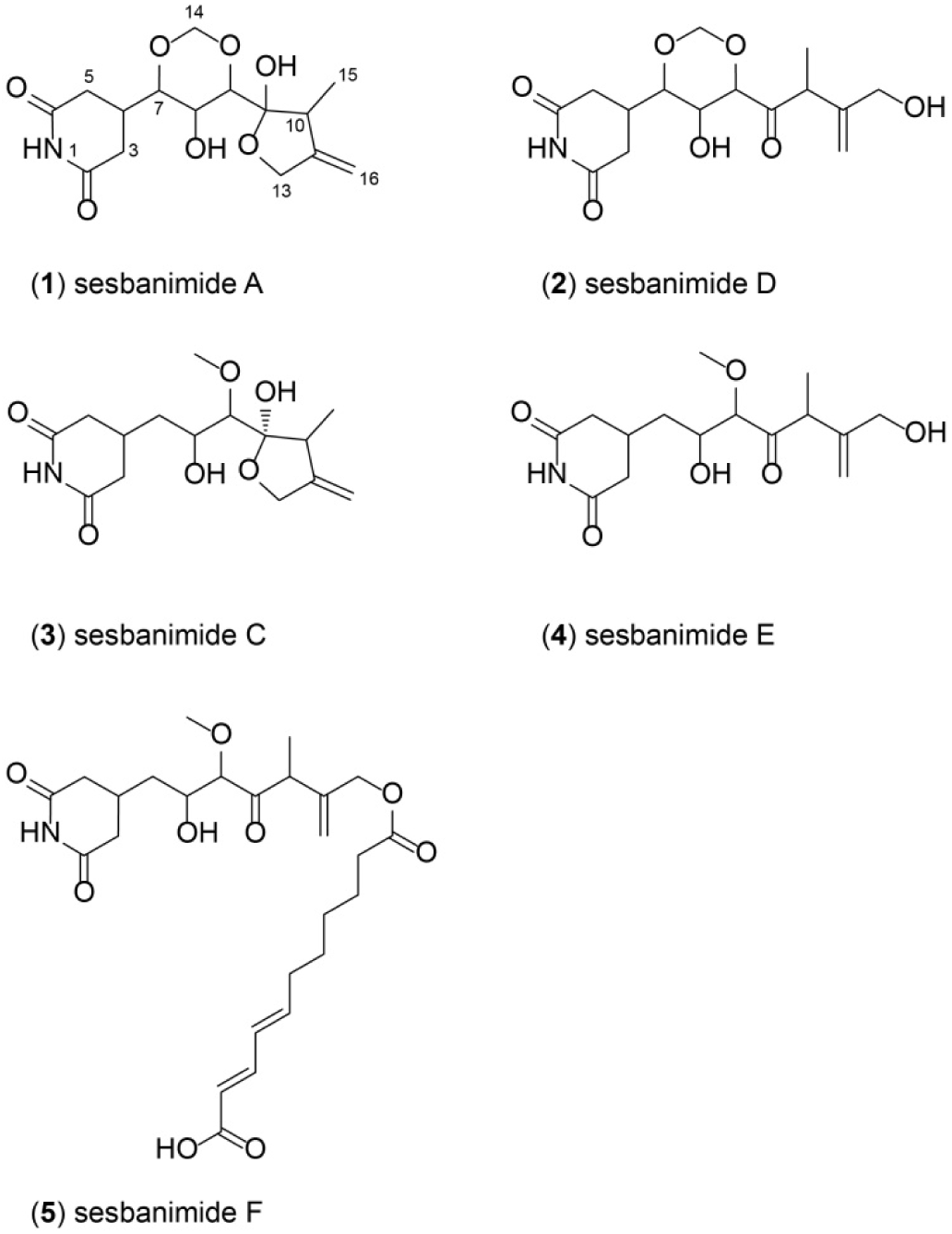
Structures of sesbanimides A (**1**) and D (**2**) isolated from PHM038 fermentation culture and sesbanimides C (**3**), D (**4**) and F (**5**) isolated from PHM037 fermentation culture.

### In vitro cytotoxicity

The bioactivity of the sesbanimide analogs was performed in cytotoxicity assays against cell lines of lung carcinoma A549 (ATCC CCL-185); colorectal carcinoma HT29 (ATCC HTB-38); and breast adenocarcinoma MDA-MB-231 (ATCC HTB-26). The cytotoxic activities in molar concentrations, as compared to control cultures, are summarized in Table 1. According to GI_50_ data, acyclic sesbanimides D and E are one to two orders of magnitude more potent than their cyclic forms A and C, respectively. Sesbanimides E and F showed similar GI_50_ results in all cell lines. Sesbanimide D is nearly one order of magnitude more potent in breast cancer cell line than in lung or colon cancer cell lines. Sesbanimide C is equally potent among all cell lines and sesbanimide A is one order of magnitude less potent in colon cancer cell line than in other ones. The most cytotoxic activity in all tested cell lines was established by sesbanimide D and the least cytotoxic activity by sesbanimide C, compared to other analogs.

**Table 1.** Cytotoxic Activity Data (Molar) of isolated sesbanimides.

| Compounds |  | Breast<br>MDA-MB-231 | Colon<br>HT29 | Lung<br>NSCLC A549 |
| --- | --- | --- | --- | --- |
| sesbanimide C | GI<sub>50</sub> | 1.6E-07 | 3.7E-07 | 2.1E-07 |
|  | TGI | 6.8E-07 | 1.1E-06 | 7.6E-07 |
|  | LC<sub>50</sub> | >3.8E-05 | >3.8E-05 | 3.5E-06 |
| sesbanimide E | GI<sub>50</sub> | 2.0E-08 | 2.3E-08 | 2.0E-08 |
|  | TGI | 1.1E-07 | 6.4E-07 | 6.4E-08 |
|  | LC<sub>50</sub> | 8.3E-06 | >3.2E-05 | 1.9E-07 |
| sesbanimide F | GI<sub>50</sub> | 2.0E-08 | 2.0E-08 | 2.0E-08 |
|  | TGI | 6.3E-06 | 1.9E-05 | 2.6E-06 |
|  | LC<sub>50</sub> | >2.0E-05 | >2.0E-05 | 7.0E-06 |
| sesbanimide A | GI<sub>50</sub> | 6.4E-07 | 1.6E-06 | 7.6E-07 |
|  | TGI | 2.5E-06 | 3.7E-06 | 2.5E-06 |
|  | LC<sub>50</sub> | 1.6E-05 | >3.1E-05 | 9.8E-06 |
| sesbanimide D | GI<sub>50</sub> | 8.6E-09 | 1.2E-08 | 1.1E-08 |
|  | TGI | 3.4E-08 | 6.1E-08 | 2.8E-08 |
|  | LC<sub>50</sub> | 1.2E-07 | >3.6E-06 | 7.9E-08 |

### Genome sequencing and identification of secondary metabolite gene clusters

The genome of *S. indica* PHM037 was assembled as a circular genome of 5,116,759 bp long with an average G+C content of 67.3 %. It comprises 4,578 coding sequences, 51 tRNAs and 2 rRNA sequences (5S-23S-16S). The genome of *L. aggregata* PHM038 was assembled in 130 contigs of 5,661,992 bp with an average G+C content of 61.5%. It comprises 5,189 coding sequences, 46 tRNAs, one 16S and one 5S rRNA. The phylogenetic analysis of 16S rRNA sequences placed the PHM037 and PHM038 strains among the genera of *Stappia* and *Labrenzia*, respectively (Fig. S25).

To explore the secondary metabolite gene clusters contained in the genomes of PHM037 and PHM038 strains we used the online platform of antiSMASH 5.0 (19). The summary of secondary metabolites gene clusters and its homologous from the MIBiG repository in PHM037 and PHM038 strains is shown in Table S1. Genome mining revealed one almost identical *trans-AT* PKS cluster in both strains. According to antiSMASH search against the MIBiG repository, genes homologous to supposed β-branching HMGS-cassette in *sbn* cluster are found in other experimentally characterized clusters: a symbiotic calyculin A biosynthetic gene cluster from uncultured Candidatus *Entotheonella* sp. (20–22); oocydin A (23); myxovirescin A1 (24); macrobrevin (25); kalimantacin A (26, 27). The protein homologues from those clusters are listed in Table S2. The more detailed analysis of the cluster (see below) showed similarities with some biosynthetic clusters of glutaramide-containing polyketides and therefore, we have proposed it as a sesbanimide biosynthetic cluster (*sbn* cluster) (Fig. 2).

**Figure 2.**
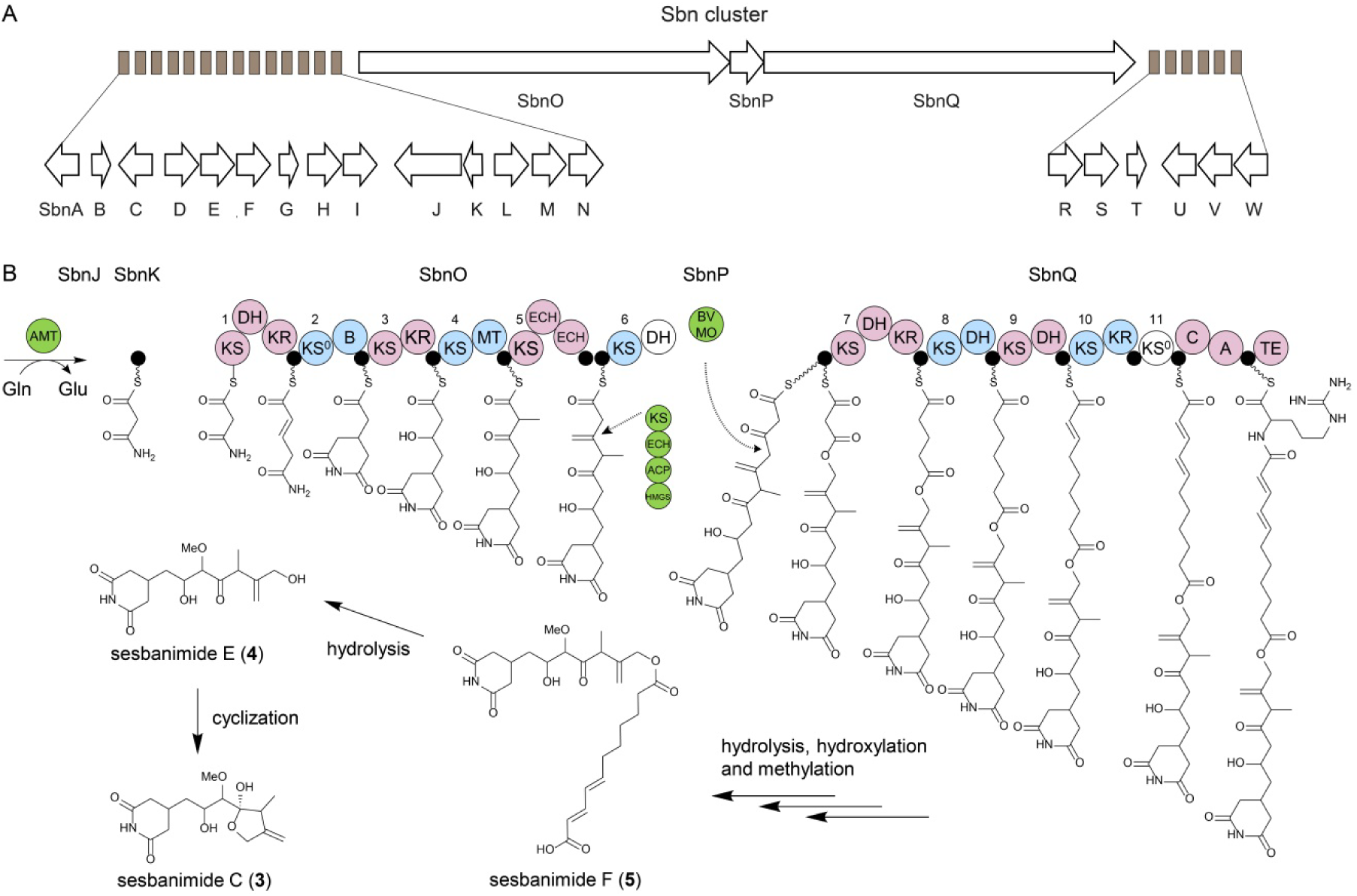
Proposed biosynthetic pathway for the assembly of sesbanimide polyketides. A) Genes comprising the *sbn* gene cluster for sesbanimide production. B) Assembly of the polyketide core structure by PKS and PKS/NRPS multimodular complexes and trans-acting components: the loading module comprising amidotranferase (AMT) and associated acyl-carrier protein (ACP); β-branching cassette comprising hydroxymethylglutaryl-CoA synthase (HMGS), stand-alone ACP, ketosynthase (KS), enoyl-CoA hydratase (ECH); Baeyer-Villiger monooxygenase (BVMO). Other PKS domains indicated are ketoreductase (KR), dehydratase (DH), branching domain (B), non-elongating ketosynthase (KS^0^), methyltransferase (MT), condensation domain (C), adenylation domain (A), peptidyl carrier protein (PCP), thioesterase domain (TE). Trans-acting proteins responsible for β-branching and oxygenation are indicated by discontinued arrows.

### Annotation of the *sbn* cluster

The *sbn* gene cluster of 64 kb encodes a modular PKS, a mixed type I PKS/non-ribosomal peptide synthetase (PKS/NRPS), 16 auxiliary enzymes with assigned functions and 5 hypothetical proteins, possibly involved in sesbanimide biosynthesis. Figure 2A shows the *sbn* cluster delimited by the marginal genes shared by PHM037 and PHM038 and named alphabetically *sbnA-W*. The *sbn* ORFs and their homologous protein are shown in Table S3. Auxiliary enzymes are *trans*-AT responsible for loading of the acyl units that serve as building blocks of polyketide assembly; *trans*-acyl hydrolase (AH) aiding the correct acyl-unit incorporation; a 4’-phosphopantetheinyl transferase (PPT) responsible for transferring the essential prosthetic group 4’-phosphopantetheine to the acyl carrier proteins (ACPs); a β-branching cassette comprising a hydroxymethylglutaryl-CoA synthase (HMGS) homolog, an ACP, a ketosynthase (KS) and an enoyl-CoA hydratase (ECH), present in other characterized polyketide clusters (28–30); a stand-alone ACP and an amidotransferase (AMT) with a glutamine hydrolyzing domain involved in glutaramide formation (6, 15); a flavin-dependent monooxygenase (OXY) homologous to a polyketide synthase component responsible for Baeyer-Villiger oxidation (31, 32); tailoring enzymes such a metallophosphoesterase (EST), a cytochrome P450, a methyltransferase (MT) (only present in PHM037 strain); and three putative ABC transport proteins.

Several putative regulators in the proximity of the *sbn* cluster were identified in both genomes. In PHM037 two ORFs coding for a putative FCD domain-containing protein with a DNA binding transcriptional factor activity were found, according to pfam, at positions 4,400,754-4,400,068 and 4,482,290-4,481,550 bp, respectively. Additionally, a TetR-like family transcriptional regulator and a sigma-54-dependent FIS-like family transcriptional regulator were found at position 4,386,957-4,387,061 bp. In PHM038 near the *sbn* cluster we have located a putative regulator at position 13,106-13,804 bp of NODE 1. Moreover, a RafR-like transcriptional regulator was found upstream of the PKS encoding gene region at position 86,332-85,217 bp of NODE 1.

### Proposed sesbanimide assembly pathway

To decipher the sesbanimide assembly pathway based on the PKS machinery, the general antiSMASH domain identification tool was complemented with the online TransATor tool (http://transator.ethz.ch) that allows for a more precise structure prediction based on clade-associated KS functions (33) (Tables S4 and S5). Subsequently, a manual inspection of the conserved motifs of the PKS and PKS/NRSPS domains was performed. In the *sbn* cluster in PHM037 strain, all KS domains in the PKS SbnO possess a conserved triad of one Cys residue and two His residues of about ~ 135 and ~175 aa downstream from the Cys site. The PKS/NRPS SbnQ have all the KS active sites besides the non-elongating KS^0^5 which is lacking a ~135 His residue. ACP domains in all modules harbor the conserved active Ser in a GxDS motif, and the ketoreductase (KR) domains feature the active site GxGxxG for NAD(P)H binding. The methyltransferase domain within SbnO has the conserved ExxxGxG motif. Out of the two dehydratase (DH) domains of SbnO, the second (C-terminal) DH does not have the conserved HxxxGxxxxP motif. Nonetheless, the pfam analysis revealed that it has a conserved C-terminal Asp194 with a N-terminal His30, common for hot-dog family proteins (34). Thioesterase (TE) domain contains both GxSxG and GxH conserved motifs.

Based on comparative analyses, computational predictions, and the elucidated chemical structures of sesbanimide intermediates we propose the sesbanimide assembly on a multidomain modular PKS and PKS/NRPS with the assistance of several *trans*-acting protein components of the enzymatic complex (Fig. 2B) based on the gene structure of PHM037. In analogy to biochemically characterized glutaramide assembly in gladiofungin molecule (17), the loading module composed of the amidotransferase SbnJ and the ACP SbnK is responsible for transferring the amino group from amino acid substrate, likely glutamine, to activated malonate. The complete glutaramide starting unit is further generated with the aid of the first 2 modules in the PKS SbnO.. The loading module together with the unusual KS-B-ACP module are present in PKSs of other glutaramides, such as cyclohexamide (15), migrastatin, iso-migrastatin, dorrigocin (36), 9-methylstreptimidone (37), and lactimidomycin (38). The remaining portion of the sesbanimide A core structure is assembled on the downstream SbnO modules. The last module contains tandem ECH domains and tandem ACPs as known from other *trans*-AT PKS modules that generate exomethylene branches (39, 40). Both ACPs possess a distinctive motif DSxxxxxW identified as important for β-branching (41). The genes *SbnF-I* encoding HMGS, ACP, KS, and ECH homologs, respectively, form a HCS cassette to assist in β-branching, a common feature of many characterized *trans*-AT PKS clusters (Table S2).

An unusual feature of the *sbn* PKS is the additional large protein SbnQ comprising four PKS and one NRPS module, which do not correspond to moieties of previously reported sesbanimides. However, the fortuitous discovery of sesbanimide F carrying an extended lipoid moiety (Fig. 2C) provides a rationale for this feature. The internal ester moiety of this congener is colinear with a split module comprising the SbnOQ interface and the monooxygenase SbnP, a module type recently shown to insert oxygen by Baeyer-Villiger oxidation resulting in ester formation (32). The lipid moiety of sesbanimide F matches well to the subsequent PKS modules, but the compound lacks the final incorporation of an amino acid suggested by the terminal NRPS module of SbnQ, hypothetically an arginine according to NRPS2 Predictor (42).

It is worth to mention that to explain the structure of the lipid moiety of sesbanimide F the two first modules of SbnQ should contain an enoyl reductase domain, but they are not present in the protein. This means that this function should be carried out by a *trans*-enoyl reductase. In addition, taking into account the non-canonical distribution of the three dehydratase and two ketoreductase domains in the four PKS modules of SbnQ, these domains should act trans-modularly coordinated to generate the double bounds present in sesbanimide F.

The presence of different sesbanimide congeners suggests that the final polyketide product might be processed into a longer intermediate sesbanimide F and shorter sesbanimides A, C, D and E, most possibly by hydrolytic esterases or amidases encoded by tailoring genes, which remains to be explored. If amide cleavage is very efficient it would explain why the fully elongated congener is not detected.

The cyclization of sesbanimide intermediates could depend on the substrate stereo-selectivity of the hydrolytic enzyme, as demonstrated by the mutational studies performed on pikromycin thioesterase (43).

Sesbanimide A and D were detected only in PHM038 strain and although their molecular structure is highly similar to C and E analogs (Fig. 1), the few structural changes could be observed. The difference in assembly of sesbanimide A on the SbnO homolog in PHM0038 strain is shown in Figure S26. Additional C7 hydroxylation is probably another tailoring modification followed by C14 cyclization by a yet unknown enzymatic process. Interestingly, in PHM038 cultures, we did not detect the ester intermediate analogous to sesbanimide F. Another difference is the absence of one ECH domain in module 5, while the strain PHM037 possesses two ECHs in tandem. Nevertheless, there is a 700 bp spacer between the AT flanking subdomain of the upstream KS and the downstream ECH in the PHM038 strain. The same domain ambiguity according to amino acid sequence is observed in various pederin family compounds (29). Finally, the downstream PKS/NRPS SbnQ is almost identical to its homologue in PHM037, with the sole exception of DH domain from the third module which conserves the HxxxGxxxxP motif.

### Comparative analysis of the *sbn* cluster with other homologous clusters

BLAST search in NCBI database lead us to detect numerous nearly identical orphan clusters among phylogenetically distant bacterial species. The genomic context of *sbn*-like clusters is shown in Figure 3. The organization of *sbnO, sbnP* and *sbnQ* genes in the *sbn* cluster was found to be identical among all analyzed bacteria. The core biosynthetic genes encoding PKSs, monooxygenase, AT and PPT are present in *S. indica* USBA 352, *Stappia* sp. BW2, *Magnetospirillum* sp. 64-120, *Stappia* sp. ARTW1, *Magnetovibrio blakemori, Roseospira navarrensis, Labrenzia alba, Phaeospirillum fulvum, Azorhizobium doebereinerae* UFLA1-100, *Rhodobium orientis* strain DSM 11290, *Labrenzia* sp. EL143, *Labrenzia* sp. Alg231-36, *Azovibrio restrictus* DSM 23866 and *Oceaniovalibus* sp. ACAM 378. Nonetheless, the number and organization of tailoring enzymes vary within species. *Magnetospirillum* sp., *A. doebereinerae* and *A. restrictus* do not contain the second AT, possessing hydrolase activity and acting as an assembly proofreading enzyme (44). Instead, those bacteria possess a type II TE coding gene in close proximity to the PKS coding genes. While all the species have one cytochrome P450 gene, some have up to four cytochrome P450 genes, annotated as biotin biosynthesis-related genes, like *Labrenzia* sp. EL143, *Labrenzia* sp. Alg231-36, *L. alba*, and *A. doebereinerae*. Interestingly, only *M. blakemori* possesses a linear amide C-N hydrolase protein, possibly involved in terminal amino acid cleavage. *P. fulvum* and *A. restrictus* do not have ABC transport proteins. *P. fulvum* has two MTs, while *A. restrictus* lacks them.

**Figure 3.**
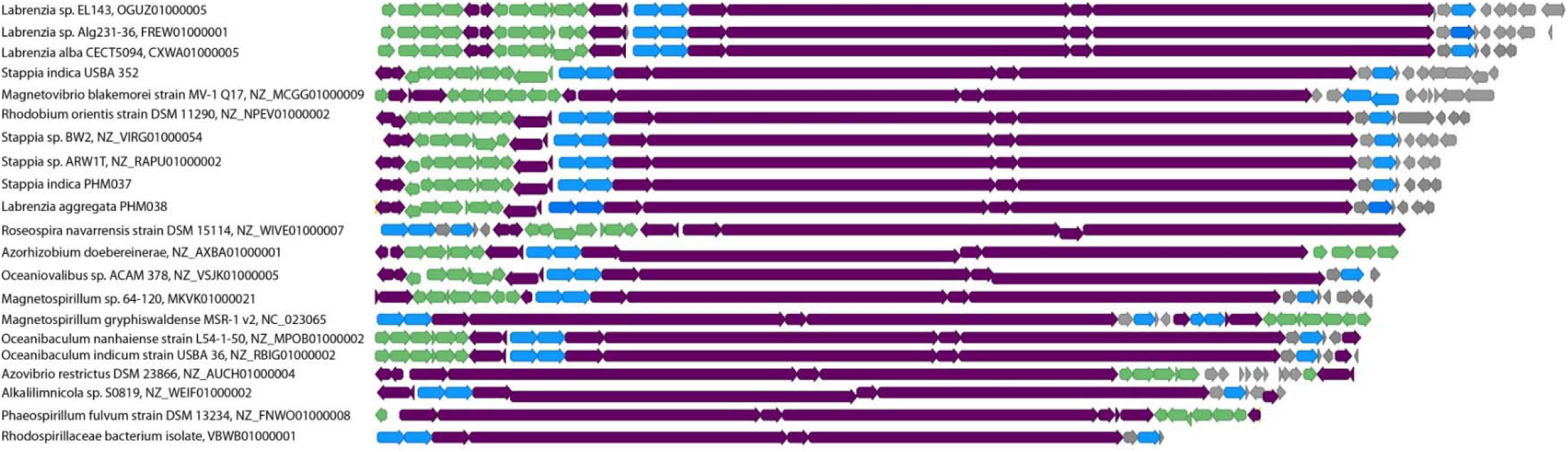
Scheme of the genomic context of the *sbn*-like clusters present in other genomes. The species and the loci accession numbers are indicated next to the cluster. The core *sbn* biosynthetic genes are shown in purple, putative ABC transport proteins in blue, tailoring enzymes in green and hypothetical proteins in grey.

Although the *sbn* clusters described here do not have transposase genes in their vicinity, *Labrenzia* sp. EL143 has a phage integrase encoding gene downstream of the cluster, *Alkalilimnicola* sp. S0819 has a transposase encoding gene upstream of the cluster and *R. navarrensis* has a transposase encoding gene downstream of the cluster. This analysis reveals an extensive horizontal gene transfer between distant organisms indicating an important ecological role of this compound.

### Comparative analysis of the modular organization of PKS and PKS/NRPS in *sbn* cluster

Surprisingly, comparative analysis of the modular architecture of SbnO revealed that the N-terminal glutaramide-forming modules, alongside with the loading module, and the C-terminal modules have originated from different gene clusters from different bacterial phyla (Fig. 4). The loading module composed of AMT and ACP proteins, denominated SbnJ and SbnK, respectively, and the first three modules of SbnO, are architecturally identical to previously identified glutaramide polyketide clusters of cyclohexamide (15), migrastatin, *iso*-migrastatin, dorrigocin (6), 9-methylstreptimidone (37) and lactimidomycin (38) as well as in glutaramide clusters of several actinomycetes and *Burkholderia* spp. genomes (14). On the other hand, the last two modules of SbnO, comprising distinctive tandem ECH and ACP domains, are absent in the glutaramide clusters from *Streptomyces* and *Burkholderia* genomes. Moreover, protein identity analysis of downstream SbnO domains revealed similarities with the pederin-family gene clusters of labrenzin (29), pederin (39, 45), nosperin (46), and cusperin (47), as well as with the clusters of onnamide (48) and oocydin A (23). Tandem ACP domains present in all genomes play an important role in β-branching, assisting the HCS cassette (49). Interestingly, a distinctive exomethylene group is present in both sesbanimide analogs, as well as in the core pyran structure of pederin family products (Fig. S27).

**Figure 4.**
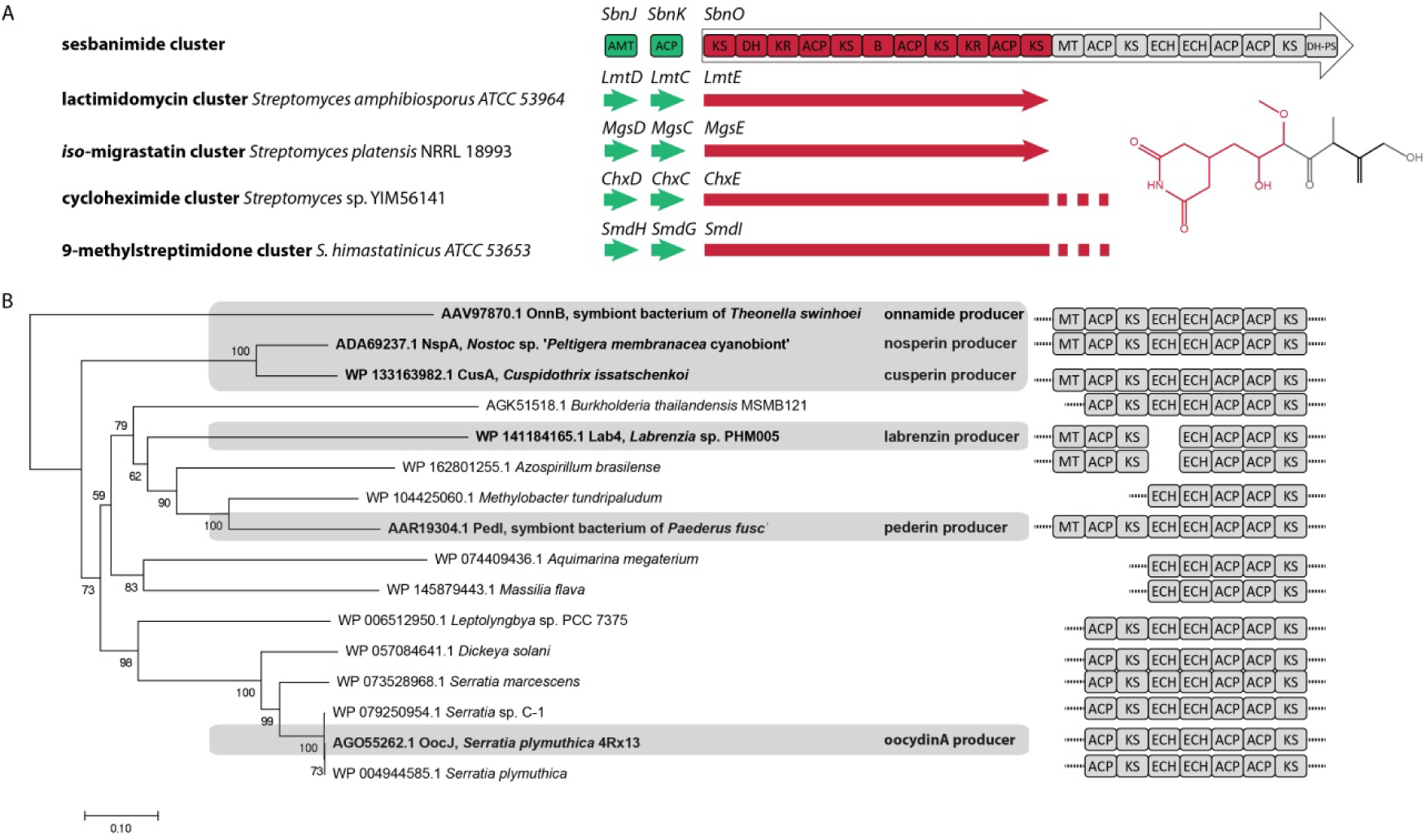
Protein identity and functionality comparisons of SbnO modules. A) Genetic scheme showing AMT, ACP and PKS from *sbn* cluster and characterized homologs in *Streptomyces* species. Sesbanimide C molecular structure (right) is shown in two colors; each color marks the moiety assembled on the respective PKS modules (the same color). B) Neighbor-joining tree based on the partial SbnO protein sequences included in the study (shadowed in grey) and the alignment with other homologous genes. Identified polyketides and their respective producers are indicated in bold. PKS domains (right) in homolog protein sequences are identified using TransATor (http://transator.ethz.ch).The scale shows evolutionary distances. Bootstrap values are indicated for each node.

Unlike SbnO, that is always present upstream SbnQ (PKS/NRPS) in *sbn*-like clusters (Fig. 3), SbnQ homologues are also present in distant PKS clusters producing structurally different polyketides lacking glutaramide moieties. To investigate the genome distribution of SbnQ, a Neighbor-joining (50) phylogenetic analysis of the closest homologues was performed (Fig. 5). Assuming OnnJ as a common ancestor from the onnamide cluster of the symbiont bacterium of *Theonella swinhoei*, we found a clade of pederin-family gene clusters producing different polyketides, *i.e*., diaphorin in a bacterium symbiont of the plant pathogen *Diaphorina citri* (51), pederin in a bacterium symbiont of the beetle *Paederus fuscipes* (52), and labrenzin in a free-living marine bacterium *Labrenzia* sp. PHM005 (29), as well as other orphan clusters of *Azospirillum brasilense* and *Methylobacter tundripaludum*. Homologues of SbnQ, like DipO, PedH and Lab13, respectively, are always present as downstream PKS/NRPS in those clusters, forming hybrid polyketide complexes with divergent upstream PKSs.

**Figure 5.**
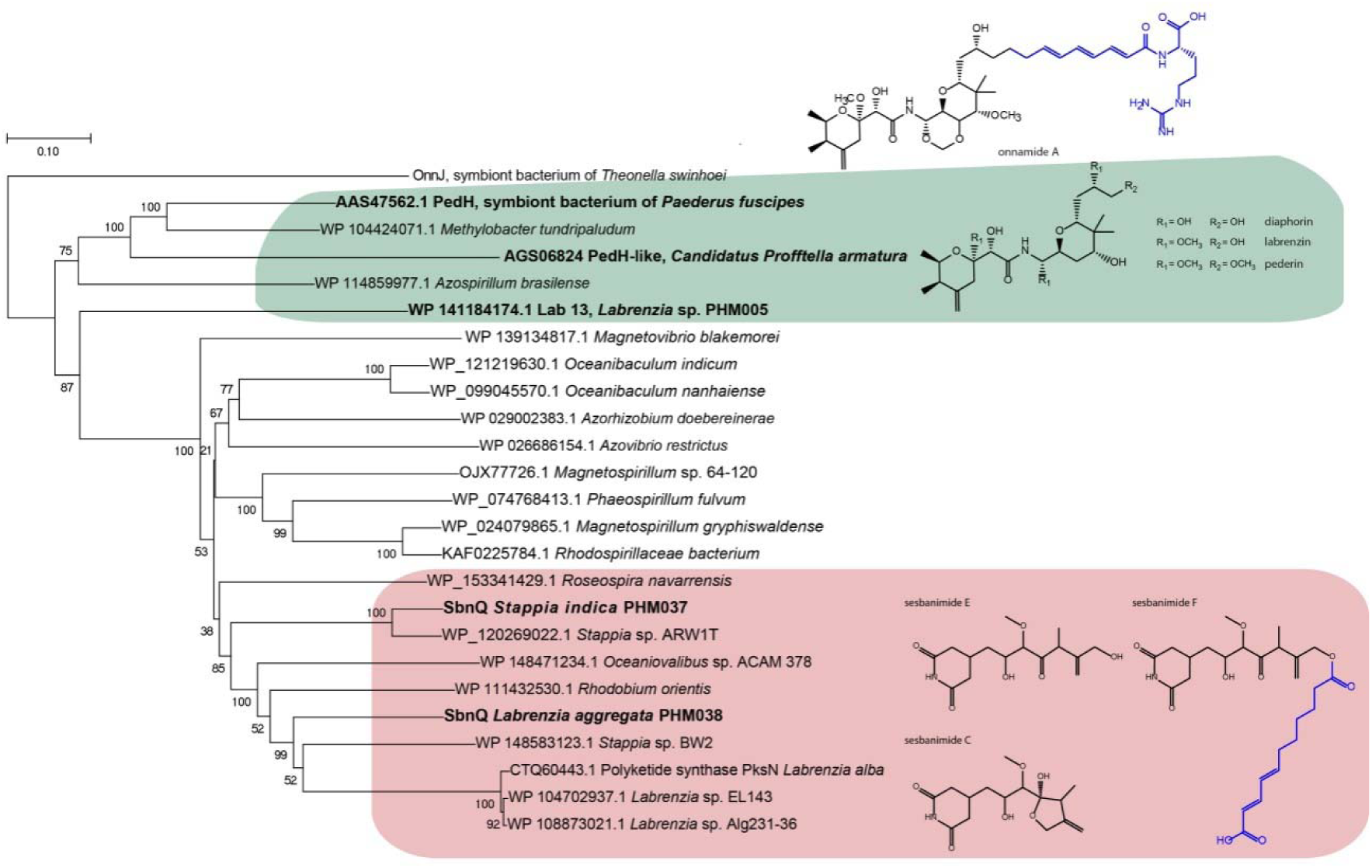
Neighbor-joining tree of SbnQ (PKS/NRPS) gene sequences, responsible for the assembly of the lipoid moiety of sesbanimide D (blue), and homologues. The characterized gene clusters are indicated in bold and their respective polyketide structures are shown (right). The scale shows evolutionary distances. Bootstrap values are indicated for each node.

Moreover, when we compared the modular architecture of SbnQ with Lab13 from the labrenzin cluster of *Labrenzia* sp. PHM005 (Fig. S28), we observed that the order of domains in the corresponding modules is nearly conserved in both PKS/NRPSs and matches the assembly of lipoid moiety of sesbanimide F in *S. indica* PHM037. Interestingly, an intermediate having a similar moiety has not been detected in the labrenzin producer yet (29). The sole modular difference detected in Lab13 is the lack of the second KS-DH-ACP module, when compared to SbnQ. Such difference implies that the polyketide moiety in the missing labrenzin analog might be shorter by one malonate building unit. The unexpected presence of almost identical downstream PKS/NRPS in unalike clusters raises important questions about the biological role of this particular molecular part, other than defense from the predators.

## Materials and Methods

### Bacteria isolation, fermentation, extraction and isolation procedures

The strain PHM037 was isolated from the sponge *Pandaros acanthifolium* collected in the Caribbean Sea (Martinique Island) in October 2003. The sample was stored at 5 °C for three days until arrived to the laboratory. Then, a suspension of the homogenized sponge in sterile sea water was spread on oligotrophic marine agar plates (ATCC Medium 357). The plates were incubated at 28 °C for one month under atmospheric pressure. The strain PHM038 was isolated from the sponge belonging to the genus of *Plakortis* in Nosy Be (Madagascar) in November 2017. The sponge was conserved in 30% glycerol at −20 °C. The frozen sample was homogenized and spread on Petri dish containing an organic medium composed of L-Asn (1.25 g/L), L-Phe (0.2 g/L), L-Pro (0.2 g/L), L-Tyr (0.2 g/L), L-Trp (0.2 g/L), glycerol (10 g/L), NaCl (5.34 g/L), KCl (0.15 g/L), MgSO_4_ (0.1 g/L), FeSO_4_ (0.1 g/L), Na_2_SO_4_ (7.5 g/L), MgCl_2_.6H_2_O (2.4 g/L), marine salts (Tropic Marin PROREEF) (10 g/L), CaCO_3_ (0.1 g/L), agar (20 g/L) and cycloheximide (200 μg/mL). The plates were incubated at 28°C during 3 weeks.

A seed culture was developed in two scale-up steps, firstly in 100 mL Erlenmeyer flasks containing 20 mL of seed medium and then 250 mL Erlenmeyer flasks with 50 mL of the same medium. The seed culture was grown on a medium containing dextrose (1.0 g/L), soluble starch (24.0 g/L), soy peptone (3.0 g/L), yeast extract (5.0 g/L), tryptone (5.0 g/L), soya flour (5.0 g/L), NaCl (5.4 g/L), KCl (0.15 g/L), MgCl_2_ (2.4 g/L), Na_2_SO_4_ (7.5 g/L), CaCO_3_ (4.0 g/L) in tap water. The strain was cultivated for 3 days at 28°C at 220 rpm. The same inoculum conditions were used for both strains.

For sebanimide production in *S. indica* PHM037, 12.5 mL of the seed medium was transferred into 2 L Erlenmeyer flasks containing 250 mL of 16B/d medium composed by brewer’s yeast (Sensient, G2025) (17.5 g/L), mannitol (76 g/L), (NH_4_)_2_SO_4_ (7 g/L), CaCO_3_ (13 g/L), FeCl_3_ (0.09 g/L) and marine salts (Tropic Marin PRO-REEF) (36 g/L). Production broth (15 L in total) was incubated for 96 h at 28 °C in a rotatory shaker at 220 rpm with 5 cm eccentricity.

For sebanimide production in *L. aggregata* PHM038, 12.5 mL of the seed medium was transferred into 2 L Erlenmeyer flasks containing 250 mL of SAM2A medium composed by glucose (10 g/L), mannitol (15 g/L), peptone (5 g/L), yeast extract (10 g/L), K_2_HPO_4_ (0.5 g/L), MgSO_4_ (0.5 g/L), FeCl_3_ (0.1 g/L), MOPS (4 g/L) and marine salts (Tropic Marin PRO-REEF) (27 g/L). Production broth (6 L in total) was incubated for 120 h at 28 °C in a rotatory shaker at 220 rpm with 5 cm eccentricity.

In both cases, after the fermentation the whole broth was centrifuged at 7000 rpm for 20 min to give aqueous supernatant which was extracted twice with EtOAc (1:1 vol:vol) and the organic phases were dried under vacuum to yield a crude extract. In this way, we obtained 649 mg of crude extract from PHM037 containing compounds **3**, **4** and **5**, and 470 mg of crude extract from PHM038 containing compounds **1** and **2**.

The extract of PHM037 was subjected to semipreparative reversed-phase HPLC using an XBridge column (19×150 mm, 5 μm) and a linear gradient of H_2_O/CH_3_CN from 5% to 50% of CH_3_CN in 30 min at a flow rate of 12.0 mL/min and UV detection at 215 nm to afford fifteen fractions (1-15). The active fraction 6 obtained between 11-13 min (30 mg) contained compounds **3** and **4**. The active fraction 13 obtained at 29 min (6 mg) contained compound **5**.

The extract of PHM038 was chromatographed on silica gel VFC (vacuum flash chromatography) system, using a stepwise gradient elution with *n*-hexane-EtOAc and EtOAc-MeOH mixtures to give eight fractions (1-8). The active fraction containing compounds **1** and **2** was eluted with EtOAc-MeOH 9:1 (25 mg) and was further purified by semi-preparative reversed-phase HPLC using a Symmetry C18 column (19×150 mm, 7 μm) and a linear gradient of H_2_O/CH_3_CN from 5% to 65% of CH_3_CN over 30 min at a flow rate of 13 mL/min, to afford fourteen fractions (1-14). The active fraction 3 between 9-10 min (5 mg) afforded a mixture of compounds **1** and **2** identified by ^1^H, ^13^C NMR and HPLCMS spectra. The active silica gel fraction eluted with EtOAc-MeOH 3:1 (28 mg) also showed by NMR spectra characteristic signals of sesbanimide compounds. This fraction was again purified by semi-preparative reversed-phase HPLC using a Symmetry C18 column (19×150 mm, 7 μm) and a linear gradient of H_2_O/CH_3_CN from 7% to 8% of CH_3_CN over 25 min, to 100% of CH_3_CN over 45 min at a flow rate of 13 mL/min, to afford eleven fractions (1-11). The active fraction 8 between 21-28 min yielded compound **1** (5 mg).

### Structure elucidation of sesbanimide analogous. NMR and mass spectra

The positive-ion ESIMS spectra were recorded using an Agilent 1100 Series LC/MS spectrometer. High Resolution Mass Spectroscopy (HRMS) was performed on an Agilent 6230 TOF LC/MS system using the ESI-MS technique. NMR spectra were recorded on a Varian “Unity 500” spectrometer at 500/125 MHz (^1^H/^13^C) and on a Varian “Unity 400” spectrometer at 400/100 MHz (^1^H/^13^C). Chemical shifts were reported in ppm using residual CD_3_OD (δ 3.31 for ^1^H and 49.0 for ^13^C), CDCl_3_ (δ 7.26 ppm for ^1^H and 77.0 ppm for ^13^C) as internal reference. 2D NMR experiments: COSY, TOCSY, HSQC and HMBC were performed using standard pulse sequences.

### Cytotoxic activity

A549 (ATCC CCL-185), HT29 (ATCC HTB-38), and MDA-MB-231 (ATCC HTB-26) cell lines were obtained from the ATCC. Cell lines were maintained in RPMI medium supplemented with 10% fetal calf serum (FCS), 2 mM L-glutamine, 100 U/mL of penicillin and 100 U/mL of streptomycin, at 37 °C and 5% CO_2_. Triplicate cultures were incubated for 72 h in the presence or absence of test compounds (at ten concentrations ranging from 10 to 0.0026 μg/mL). For quantitative estimation of cytotoxicity, the colorimetric sulforhodamine B (SRB) method was used (53). Briefly, cells were washed twice with PBS, fixed for 15 min in 1% glutaraldehyde solution, rinsed twice in PBS, and stained in 0.4% SRB solution for 30 min at room temperature. Cells were then rinsed several times with 1% acetic acid solution and air-dried. Sulforhodamine B was then extracted in 10 mM Trizma base solution and the absorbance measured at 490 nm.

Using the mean+SD of triplicate cultures, a dose-response curve was automatically generated using nonlinear regression analysis. Three reference parameters were calculated (NCI algorithm) by automatic interpolation: GI_50_ (compound concentration that produces 50% cell growth inhibition, as compared to control cultures); TGI (total cell growth inhibition (cytostatic effect), as compared to control cultures), and LC_50_ (compound concentration that produces 50% net cell killing (cytotoxic effect).

### Genome sequencing, assembly and annotation

Overnight cultures of PHM037 and PHM038 were used for the genomic DNA extraction using Blood & Cell Culture DNA Mini Kit (Qiagen). The genome of *S. indica* PHM037 was sequenced using PacBio RSII sequencer and assembled by Macrogen (http://www.macrogen.com) into a single contig. It was re-sequenced by Illumina HiSeq technology using a 250 bp paired end protocol provided by MicrobesNG http://www.microbesng.uk). The Illumina reads were mapped to PHM037 genome sequenced by PacBio using the Geneious v.10.0.2 software to manually correct PacBio sequencing errors. The sequencing and the assembly of the genome of PHM038 was performed by MicrobesNG (http://www.microbesng.uk). The genomes of *S. indica* PHM037 and *L. aggregata* PHM038 were annotated using the NCBI Prokaryotic Genome Annotation Pipeline (PGAP). This Whole Genome Shotgun project has been deposited at DDBJ/ENA/GenBank under the accessions CP046908 and JABFCZ000000000 for strains PHM037 and PHM038, respectively. The versions described in this paper are versions CP046908 and JABFCZ010000000. The SRA data was submitted under accessions SRR10686572 and SRR10686571 for PHM037 strain and SUB7382819 for PHM038 strain. The sesbanimide gene cluster was submitted to MIBiG repository under accession number BGC0002082. Genome mining for secondary metabolites gene clusters was done by automated antibiotics and Secondary Metabolite Analysis SHell (antiSMASH) 5.0 (19). Functional assignation to *trans*-AT PKS cluster ORFs was based on BLASTp searches. Additionally, PKS and NRPS domain architecture and specificities were consulted using TransATor (http://transator.ethz.ch) and pfam database. Routine bioinformatic analyses were performed by Geneious Prime version 2020.0.4 created by Biomatters (http://www.geneious.com).

### Phylogenetic analysis

Partial nucleotide sequences of the 16S rDNA gene were aligned with CLUSTAL X v2 using the default parameters, and verified manually in order to maximize positional homology. All positions with gaps were omitted for the phylogenetic reconstructions. Additionally, sequences retrieved from SILVA Refseq database (www.arb-silva.de), were also included in the analyses. The data set was subjected to the NJ and maximum likelihood (ML) method of phylogenetic inference. Analyses were carried out with PAUP v4.0b10 (54). The Akaike information criterion (AIC) implemented in Modeltest v2 (55) was used to select the evolutionary model that best fit the empirical data set. The TIM (“transitional model”) was selected as the best-fit model to the dataset. Robustness of the resulting tree was tested with 1000 bootstrapping (56). The PKS and PKS/NRPS sequences were collected from the public databases. The sequences were aligned by Clustal Omega build in Geneious Prime (http://www.geneious.com). The NJ analysis was performed using MEGA7 (57). For tree resampling a bootstrap of 1000 replicates was used. The evolutionary distances were computed using the Poisson correction method (58).

## Author contributions

D.K. conceived the idea, developed the theory, performed the bioinformatics analyses and wrote the first draft of the manuscript. L.C. performed purifications, structure elucidation of compounds and wrote the manuscript chemical part. P.R. performed bacterial cultures. E.G. performed phylogenetic analysis of 16S rDNA. C.S. selected the strains for genomic mining. J.P. and S.L.-M. contributed to develop the biosynthetic model and to discuss the functions and comparisons of gene clusters. F.C., C.C., B.G. and J.L.G. were in charge of overall direction and planning. All authors provided critical feedback and helped shape the research, analysis and manuscript.

The authors declare no competing interest.

## Acknowledgements

The present study was funded by the Ministry of Economy and Competitiveness of Spain under the program RETOS-COLABORACIÓN with the project number RTC-2016-4892-1 (DESPOL). The authors would like to thank Cristina Lillo for performing the MS experiments and Ana Valencia for technical assistance.

## References

1. R. G. Powell, R. D. Plattner, M. Suffness, Occurrence of sesbanimide in seeds of toxic sesbania species. Weed Sci. 38, 148–152 (1990).

2. R. G. Powell, et al., Sesbanimide, a potent ntitumor substance from *Sesbania drummondii* seed. J. Am. Chem. Soc. 105, 3739–3741 (1983).

3. R. G. Powell, C. R. Smith, An investigation of the antitumor activity of *Sesbania drummondii*. J. Nat. Prod. 44, 86–90 (1981).

4. R. G. Powel, C. R. Smith, R. V Madrigal, Antitumor activity of *Sebania versicara, S. punicea* and *S. drummondii*. Planta Med. 30, 1–8 (1976).

5. C. P. Gorst-Allman, P. S. Steyn, R. Vleggaar, N. Grobbelaar, Structure elucidation of sesbanimide using high-field n.m.r. spectroscopy. J. Chem. Soc. Perkin Trans. 1, 1311–1314 (1984).

6. S. K. Lim, et al., Iso-migrastatin, migrastatin, and dorrigocin production in *Streptomyces platensis* NRRL 18993 is governed by a single biosynthetic machinery featuring an acyltransferase-less type I polyketide synthase. J. Biol. Chem. 284, 29746–29756 (2009).

7. J. Ju, et al., Lactimidomycin, iso-migrastatin and related glutarimide-containing 12-membered macrolides are extremely potent inhibitors of cell migration. J. Am. Chem. Soc. 131, 1370–1371 (2009).

8. S. R. Rajski, B. Shen, Multifaceted modes of action for the glutarimide-containing polyketides revealed. ChemBioChem 11, 1951–1954 (2010).

9. S. Huang, et al., Cycloheximide and congeners as inhibitors of eukaryotic protein synthesis from endophytic actinomycetes *Streptomyces* sps. YIM56132 and. J. Antibiot. (Tokyo)., 163–166 (2011).

10. T. Schneider-Poetsch, et al., Inhibition of eukaryotic translation elongation by cycloheximide and lactimidomycin. Nat. Chem. Biol. 6, 209–217 (2010).

11. N. Saito, F. Suzuki, K. Sasaki, N. Ishida, Antiviral and interferon inducing activity of a new glutarimide antibiotic, 9 methylstreptimidone. Antimicrob. Agents Chemother. 10, 14–19 (1976).

12. X. L. Zhao, et al., Two new glutarimide antibiotics from *Streptomyces* sp. HS-NF-780. J. Antibiot. (Tokyo). 72, 241–245 (2019).

13. D. Zhang, W. Yi, H. Ge, Z. Zhang, B. Wu, Bioactive streptoglutarimides A-J from the marine-derived *Streptomyces* sp. ZZ741. J. Nat. Prod. 82, 2800–2808 (2019).

14. E. R. Stulberg, et al., Genomic and secondary metabolite analyses of *Streptomyces* sp. 2AW provide insight into the evolution of the cycloheximide pathway. Front. Microbiol. 7, 1–12 (2016).

15. M. Yin, et al., Cycloheximide and actiphenol production in streptomyces sp. YIM56141 governed by single biosynthetic machinery featuring an acyltransferase-less type i polyketide synthase. Org. Lett. 16, 3072–3075 (2014).

16. K. V. Rao, C-73: A Metabolic product of *Streptomyces albulus*. J. Org. Chem. 25, 661–662 (1960).

17. S. P. Niehs, et al., Insect-associated bacteria assemble the antifungal butenolide gladiofungin by non-canonical polyketide chain termination. Angew. Chemie Int. Ed. (2020) https://doi.org/10.1002/anie.202005711.

18. C. Acebal, et al., Two marine agrobacterium producers of sesbanimide antibiotics. J. Antibiot. (Tokyo). 51, 64–67 (1998).

19. K. Blin, et al., antiSMASH 5.0: updates to the secondary metabolite genome mining pipeline. Nucleic Acids Res. 47, W81–W87 (2019).

20. T. Wakimoto, et al., Calyculin biogenesis from a pyrophosphate protoxin produced by a sponge symbiont. Nat. Chem. Biol. 10, 648–655 (2014).

21. A. E. Fagerholm, D. Habrant, A. M. P. Koskinen, Calyculins and related marine natural products as serine-threonine protein phosphatase PP1 and PP2A inhibitors and total syntheses of calyculin A, B, and C. Mar. Drugs 8, 122–172 (2010).

22. A. R. Uria, Capturing natural product biosynthetic pathways from uncultivated symbiotic bacteria of marine sponges through metagenome mining: A mini-review. SQUALEN, Bull. Mar. Fish. Postharvest Biotechnol. 10, 35–49 (2015).

23. M. A. Matilla, H. Stöckmann, F. J. Leeper, G. P. C. Salmond, Bacterial biosynthetic gene clusters encoding the anti-cancer haterumalide class of molecules: Biogenesis of the broad spectrum antifungal and anti-oomycete compound, oocydin A. J. Biol. Chem. 287, 39125–39138 (2012).

24. V. Simunovic, et al., Myxovirescin A biosynthesis is directed by hybrid polyketide synthases/nonribosomal peptide synthetase, 3-hydroxy-3-methylglutaryl-CoA synthases, and trans-acting acyltransferases. ChemBioChem 7, 1206–1220 (2006).

25. Y. Bai, et al., Functional overlap of the Arabidopsis leaf and root microbiota. Nature 528, 364–369 (2015).

26. W. Mattheus, et al., Isolation and purification of a new kalimantacin / batumin-related polyketide antibiotic and slucidation of its biosynthesis gene cluster. Chem. Biol. 17, 149–159 (2010).

27. B. Uytterhoeven, et al., Systematic analysis of the kalimantacin assembly line NRPS module using an adapted targeted mutagenesis approach. Microbiologyopen 5, 279–286 (2016).

28. T. J. Buchholz, et al., Polyketide β-branching in bryostatin biosynthesis: Identification of surrogate acetyl-ACP donors for BryR, an HMG-ACP synthase. Chem. Biol. 17, 1092–1100 (2010).

29. D. Kačar, et al., Genome of Labrenzia sp. PHM005 Reveals a complete and active trans-AT PKS gene cluster for the biosynthesis of labrenzin. 10, 1–14 (2019).

30. L. Gu, et al., Metamorphic enzyme assembly in polyketide diversification. Nature 459, 731–735 (2009).

31. M. C. Tang, H. Y. He, F. Zhang, G. L. Tang, Baeyer-villiger oxidation of acyl carrier protein-tethered thioester to acyl carrier protein-linked thiocarbonate catalyzed by a monooxygenase domain in FR901464 biosynthesis. ACS Catal. 3, 444–447 (2013).

32. R. A. Meoded, et al., A polyketide synthase component for oxygen insertion into polyketide backbones. Angew. Chemie - Int. Ed. 57, 11644–11648 (2018).

33. E. J. N. Helfrich, et al., Automated structure prediction of trans-acyltransferase polyketide synthase products. Nat. Chem. Biol. 15, 813–821 (2019).

34. S. C. Dillon, A. Bateman, The Hotdog fold: Wrapping up a superfamily of thioesterases and dehydratases. BMC Bioinformatics 5, 1–14 (2004).

35. S. Sundaram, D. Heine, C. Hertweck, Polyketide synthase chimeras reveal key role of ketosynthase domain in chain branching. Nat. Chem. Biol. 11, 949–951 (2015).

36. S.-K. Lim, et al., *iso*-Migrastatin, migrastatin, and dorrigocin production in *Streptomyces platensis* NRRL 18993 is governed by a single biosynthetic machinery featuring an acyltransferase-less type I polyketide synthase. J. Biol. Chem. 284, 29746–29756 (2009).

37. B. Wang, et al., Biosynthesis of 9⍰methylstreptimidone involves a new decarboxylative step for polyketide terminal diene formation. 7–10 (2013).

38. J. W. Seo, et al., Comparative characterization of the lactimidomycin and iso-migrastatin biosynthetic machineries revealing unusual features for acyltransferase-less type I polyketide synthases and providing an opportunity to engineer new analogues. Biochemistry 53, 7854–7865 (2014).

39. J. Piel, A polyketide synthase-peptide synthetase gene cluster from an uncultured bacterial symbiont of Paederus beetles. Proc Natl Acad Sci U S A 99, 14002–14007 (2002).

40. E. J. N. Helfrich, J. Piel, Biosynthesis of polyketides by trans-AT polyketide synthases. Nat. Prod. Rep. 33, 231–316 (2016).

41. S. Kosol, M. Jenner, J. R. Lewandowski, G. L. Challis, Protein–protein interactions in *trans*-AT polyketide synthases. Nat. Prod. Rep., 1097–1109 (2018).

42. M. Röttig, et al., NRPSpredictor2 - A web server for predicting NRPS adenylation domain specificity. Nucleic Acids Res. 39, 362–367 (2011).

43. A. A. Koch, et al., A single active site mutation in the pikromycin thioesterase generates a more effective macrocyclization catalyst. J. Am. Chem. Soc. 139, 13456–13465 (2017).

44. N. Dimitrova, et al., Polyketide proofreading by an acyltransferase-like enzyme. PLoS One 32, 736–740 (2017).

45. J. Piel, G. Wen, M. Platzer, D. Hui, Unprecedented diversity of catalytic domains in the first four modules of the putative pederin polyketide synthase. ChemBioChem 5, 93–98 (2004).

46. A. Kampa, et al., Metagenomic natural product discovery in lichen provides evidence for a family of biosynthetic pathways in diverse symbioses. Proc. Natl. Acad. Sci. 110, E3129–E3137 (2013).

47. A. Kust, et al., Discovery of a pederin family compound in a nonsymbiotic bloom-forming *Cyanobacterium*. ACS Chem. Biol. 13, 1123–1129 (2018).

48. J. Piel, et al., Antitumor polyketide biosynthesis by an uncultivated bacterial symbiont of the marine sponge *Theonella swinhoei*. PNAS 101, 16222–16227 (2004).

49. A. S. Haines, et al., A conserved motif flags acyl carrier proteins for β-branching in polyketide synthesis. Nat. Chem. Biol. 9, 685–692 (2013).

50. N. Saitou, M. Nei, The neighbor-joining method: a new method for reconstructing phylogenetic trees. Mol. Biol. Evol. 4, 406–425 (1987).

51. A. Nakabachi, et al., Defensive bacteriome symbiont with a drastically reduced genome. Curr. Biol. 23, 1478–1484 (2013).

52. J. Piel, D. Hui, N. Fusetani, S. Matsunaga, Targeting modular polyketide synthases with iteratively acting acyltransferases from metagenomes of uncultured bacterial consortia. Environ. Microbiol. 6, 921–927 (2004).

53. K. T. Papazisis, G. D. Geromichalos, K. A. Dimitriadis, A. H. Kortsaris, Optimization of the sulforhodamine B colorimetric assay. J. Immunol. Methods 208, 151–158 (1997).

54. D. L. Swofford, Phylogenetic analysis using parsimony (paup), version 4. Sunderland, MA Sinauer Assoc. (1998).

55. D. Posada, K. A. Crandall, MODELTEST: testing the model of DNA substitution. Bioinformatics 14, 817–818 (1998).

56. J. Felsenstein, Confidence limits on phylogenies: An approach using the bootstrap. Evolution (N. Y). 39, 783 (1985).

57. S. Kumar, G. Stecher, K. Tamura, MEGA7: Molecular evolutionary genetics analysis version 7.0 for bigger datasets. Mol. Biol. Evol. 33, 1870–1874 (2016).

58. E. Zuckerkandl, L. Pauling, Evolutionary divergence and convergence in proteins. Evol. Genes Proteins, 97–166 (1965).

