## Supplementary material for "Identification of trans-AT polyketide clusters in two marine bacteria reveals cryptic similarities between distinct symbiosis factors"

**This PDF file includes:**

Figures S1 to S28

Tables S1 to S5

Index

### Supplementary information on chemical elucidation of the compounds

#### Figures S1-S4.1D and 2D NMR spectra of 1in CDCl_3_

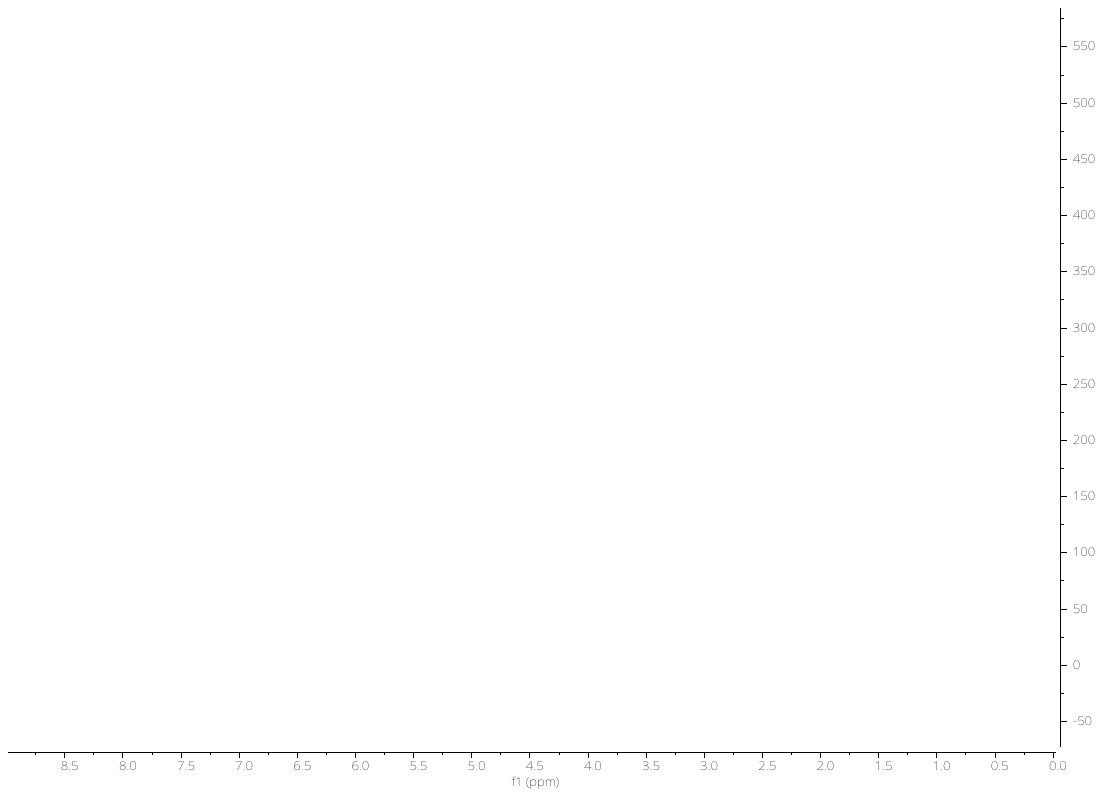

**Fig. S1.**^1^H NMR (500 MHz) Spectrum of Compound **1** (Sesbanimide A) in CDCl_3_

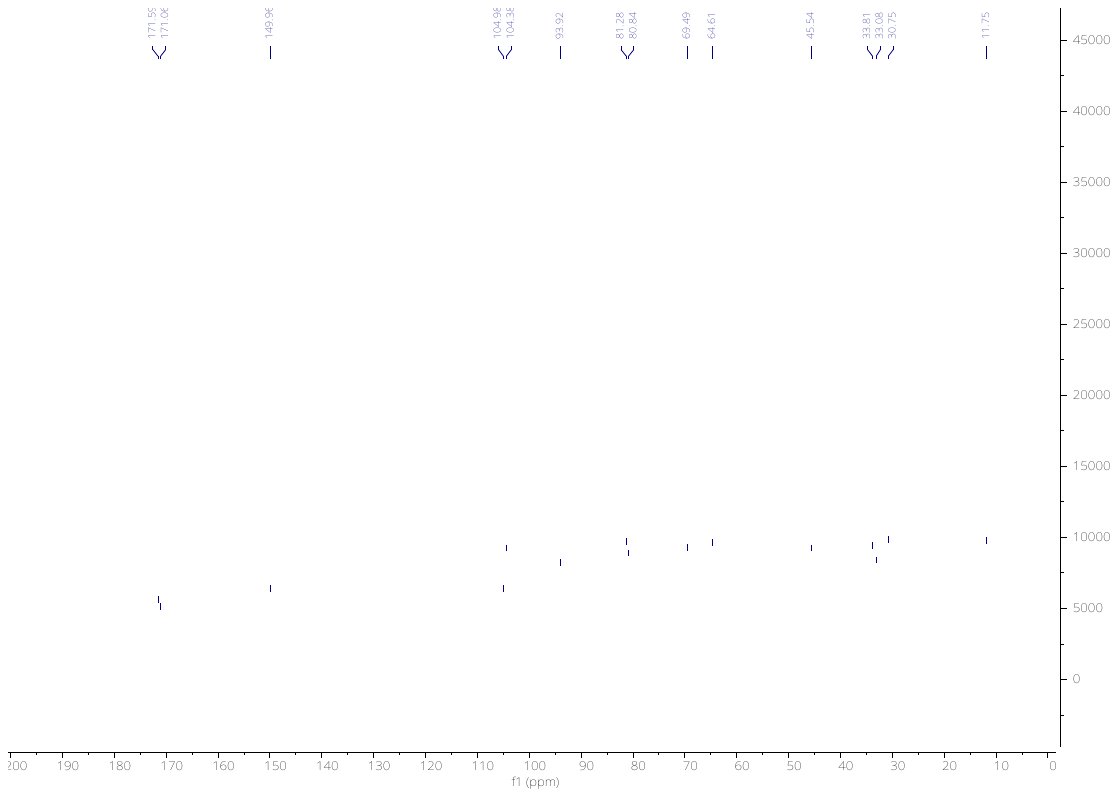

**Fig. S2.**^13^C NMR (100 MHz) Spectrum of Compound **1** in CDCl_3_

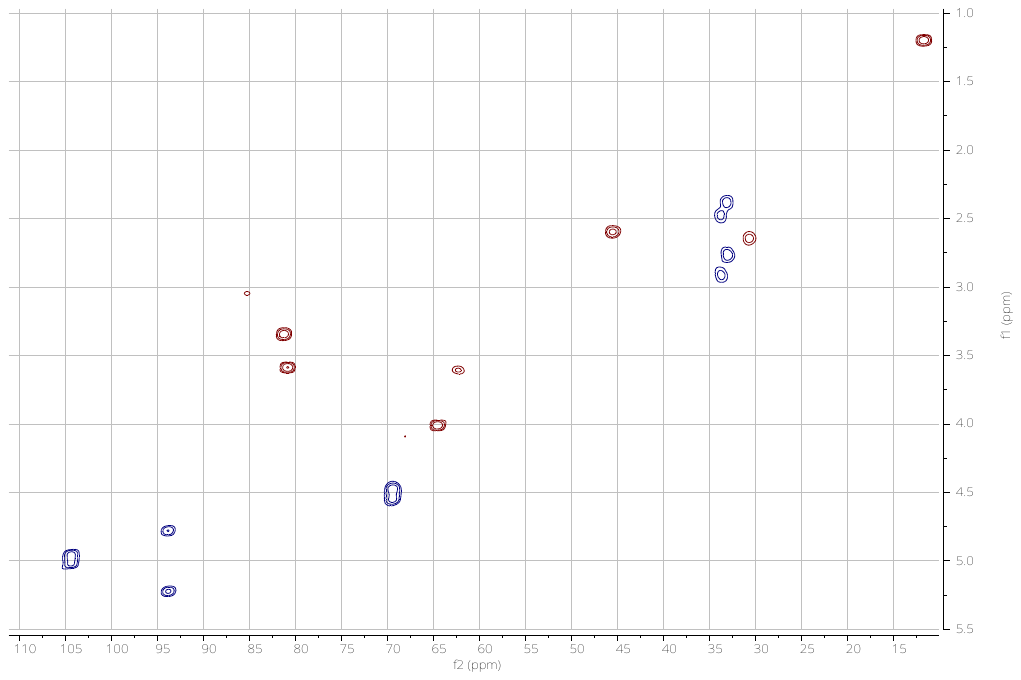

**Fig. S3.** gHSQC (400 MHz) Spectrum of Compound **1** in CDCl_3_

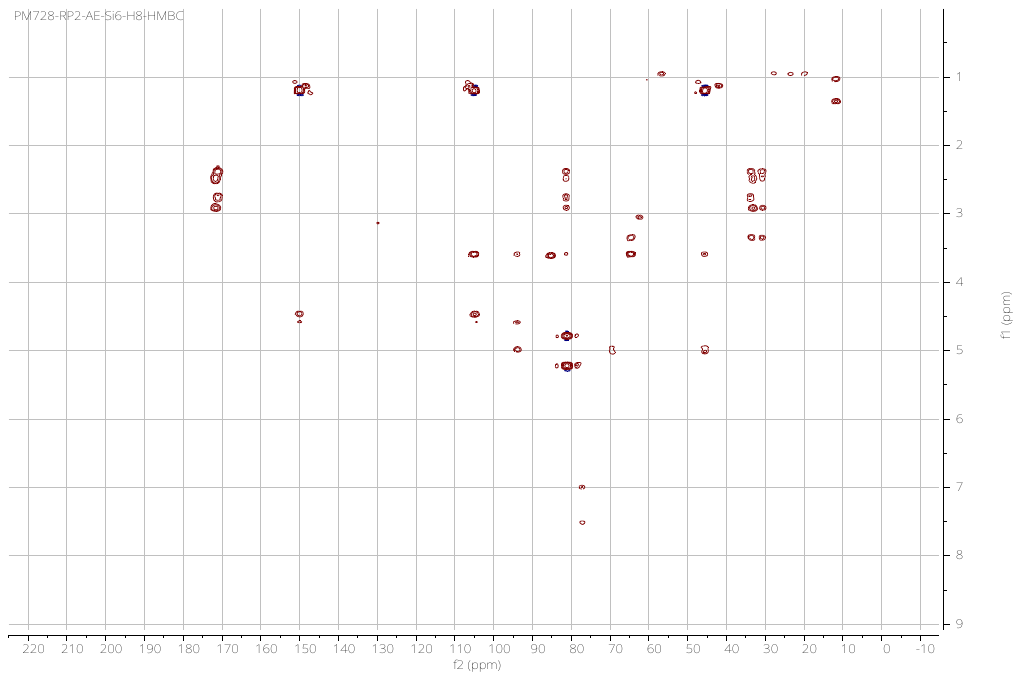

**Fig. S4.** gHMBC (400 MHz) Spectrum of Compound **1** in CDCl_3_

#### Figures S5-S9.1D and 2D NMR spectra of 2 in CD_3_OD

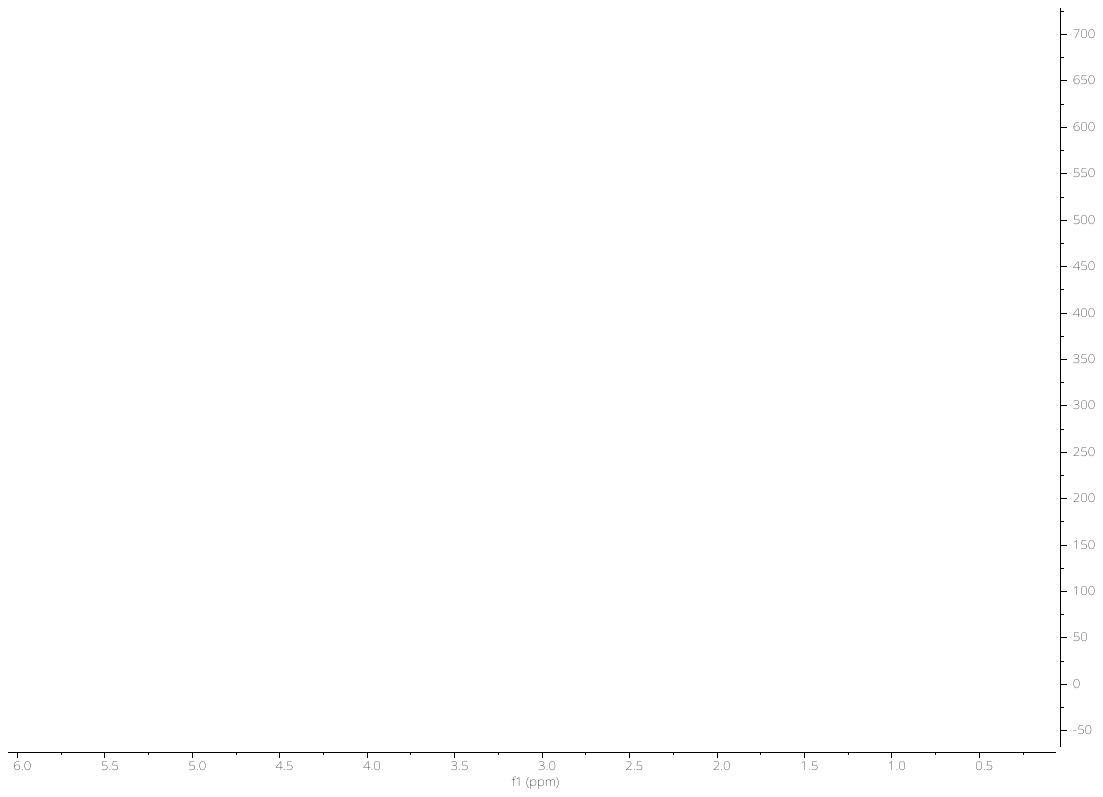

**Fig. S5.**^1^H NMR (400 MHz) Spectrum of Compound **2** in CD_3_OD

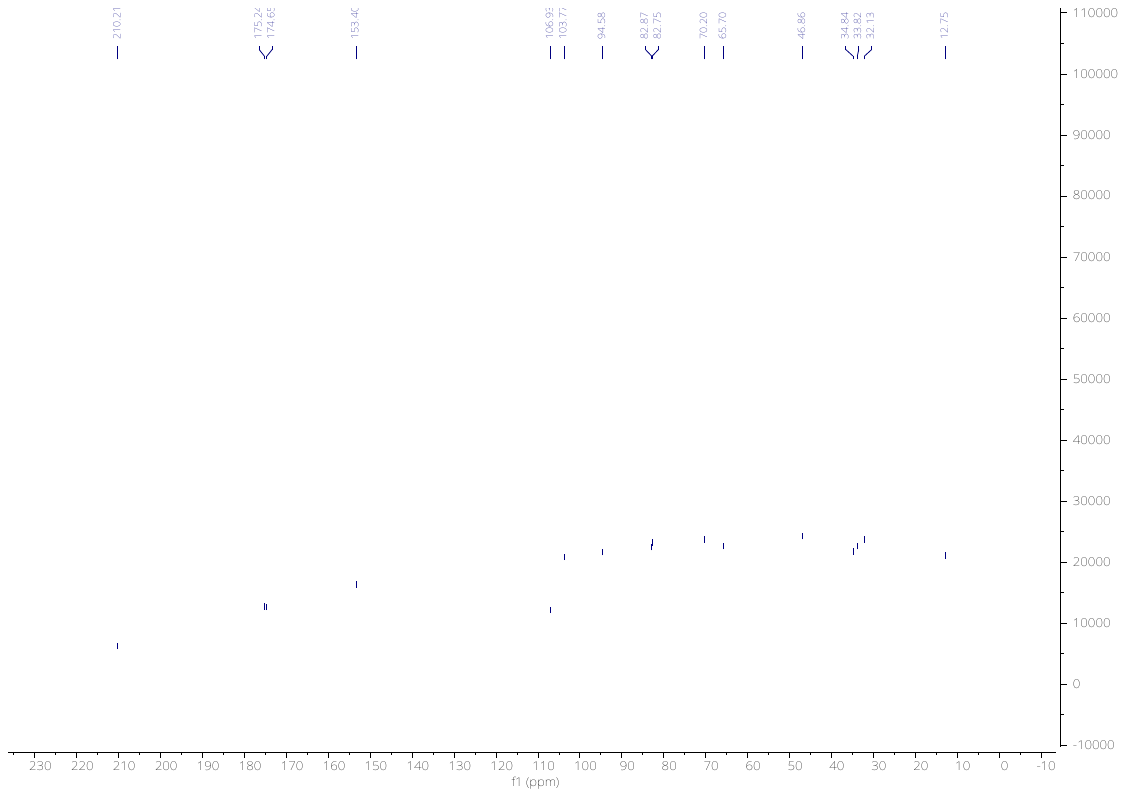

**Fig. S6.**^13^C NMR (400 MHz) Spectrum of Compound **2** in CD_3_OD

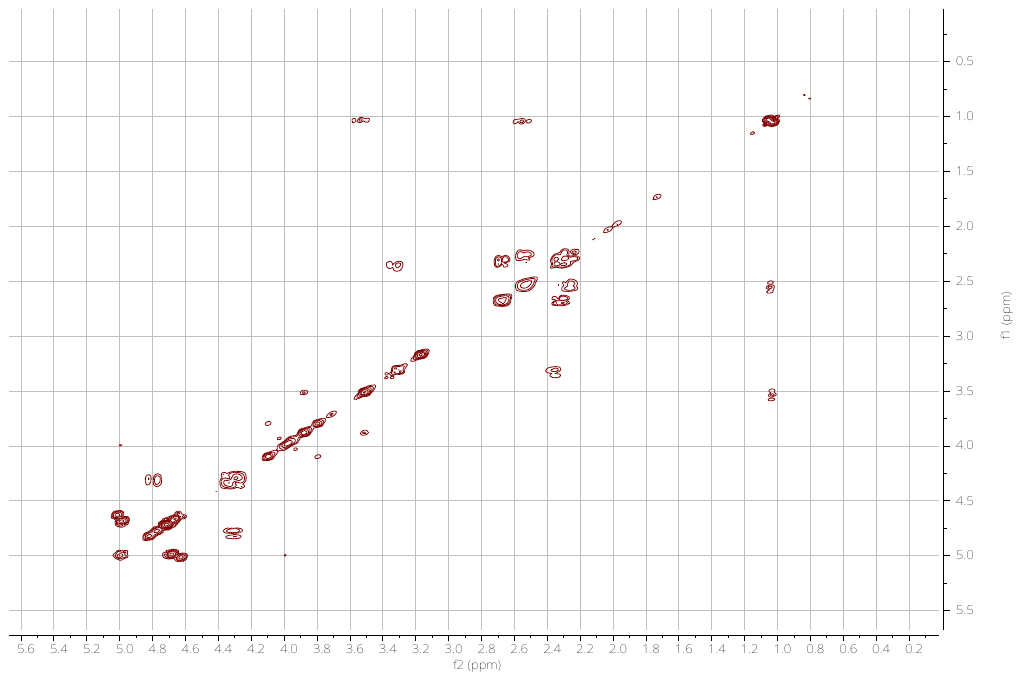

**Fig. S7.** gCOSY (400 MHz) Spectrum of Compound **2** in CD_3_OD

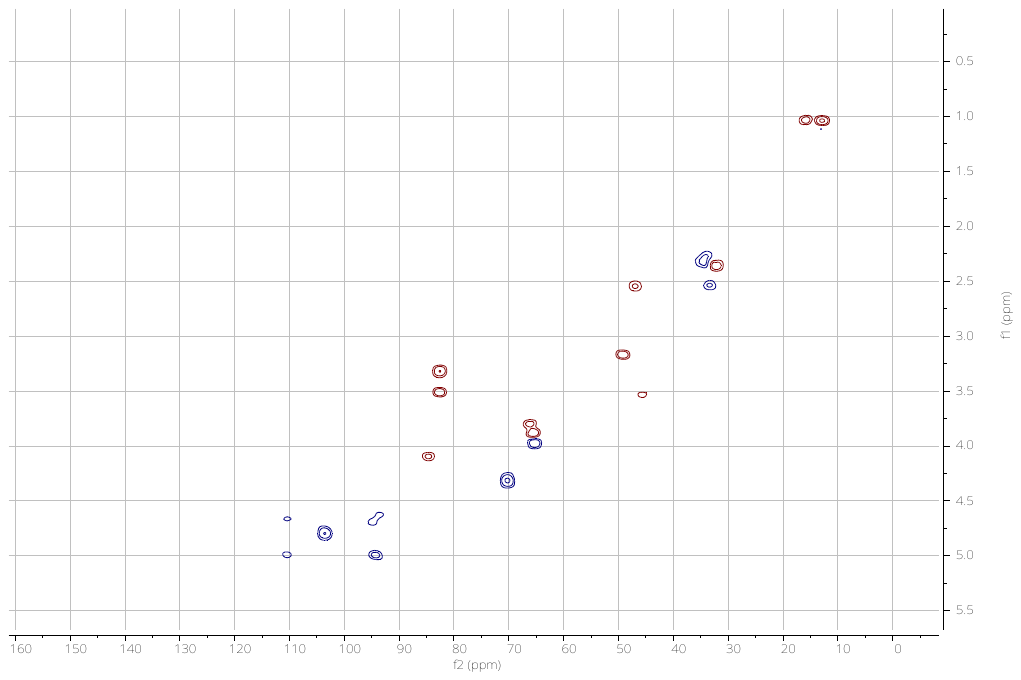

**Fig. S8.** gHSQC (400 MHz) Spectrum of Compound **2** in CDCl_3_

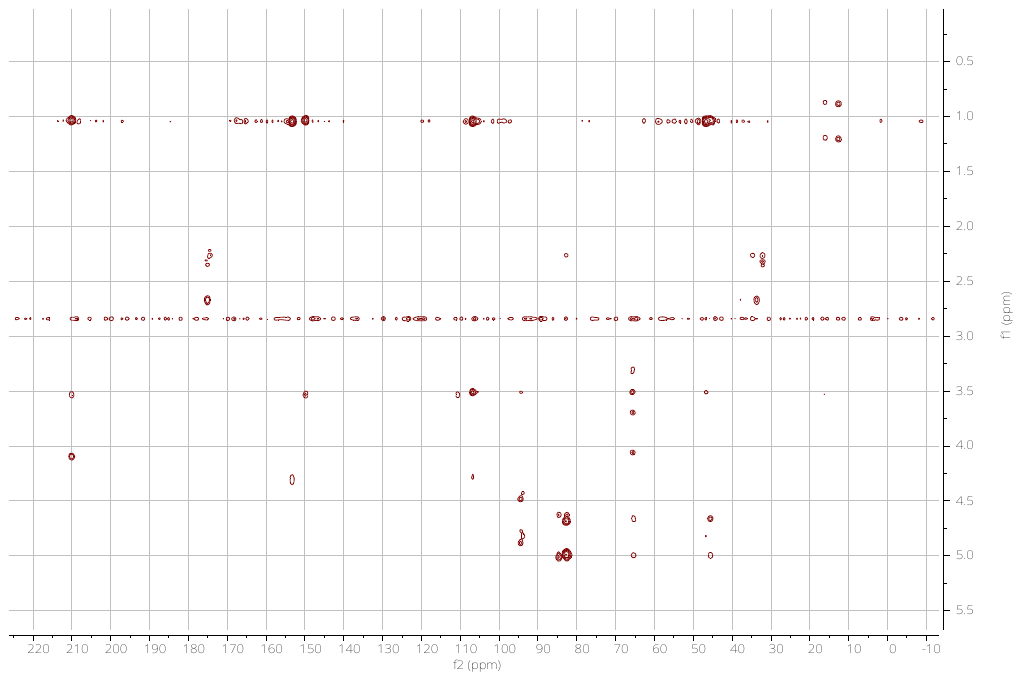

**Fig. S9.** gHMBC (400 MHz) Spectrum of Compound **2** in CD_3_OD

#### Figures S10-S11.1D NMR spectra of 3 in CD_3_OD

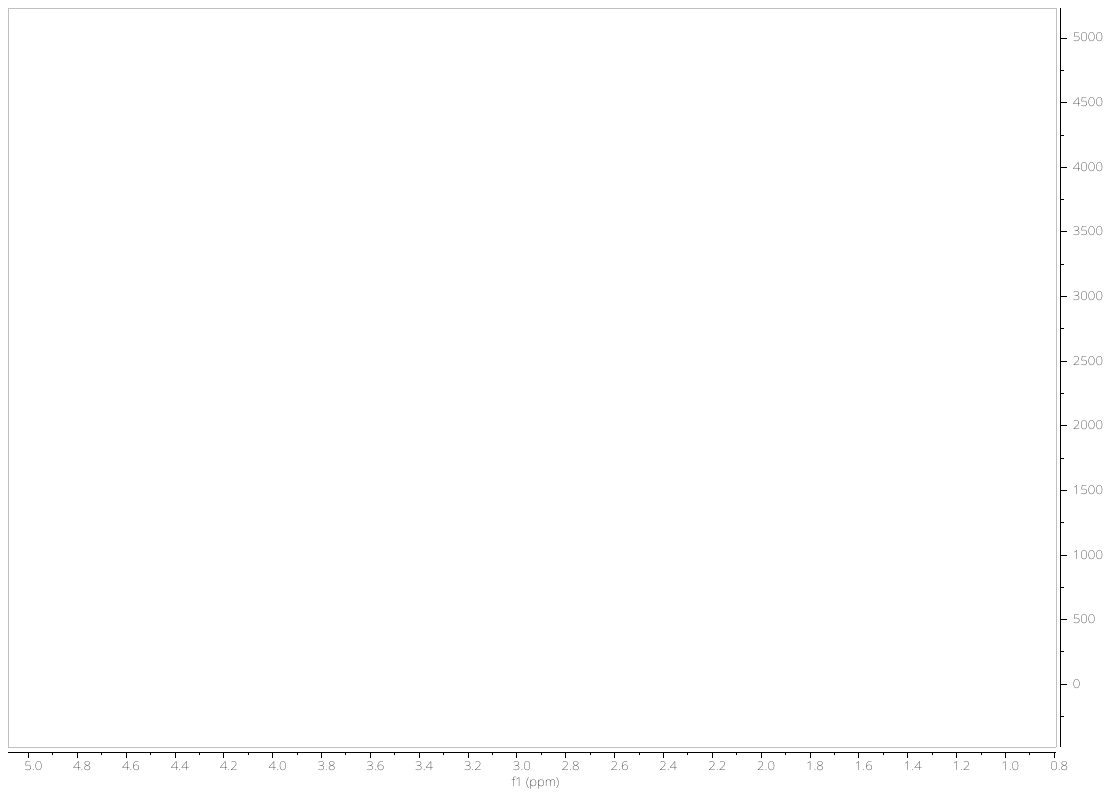

**Fig. S10.**^1^H NMR (400 MHz) Spectrum of Compound **3** in CD_3_OD

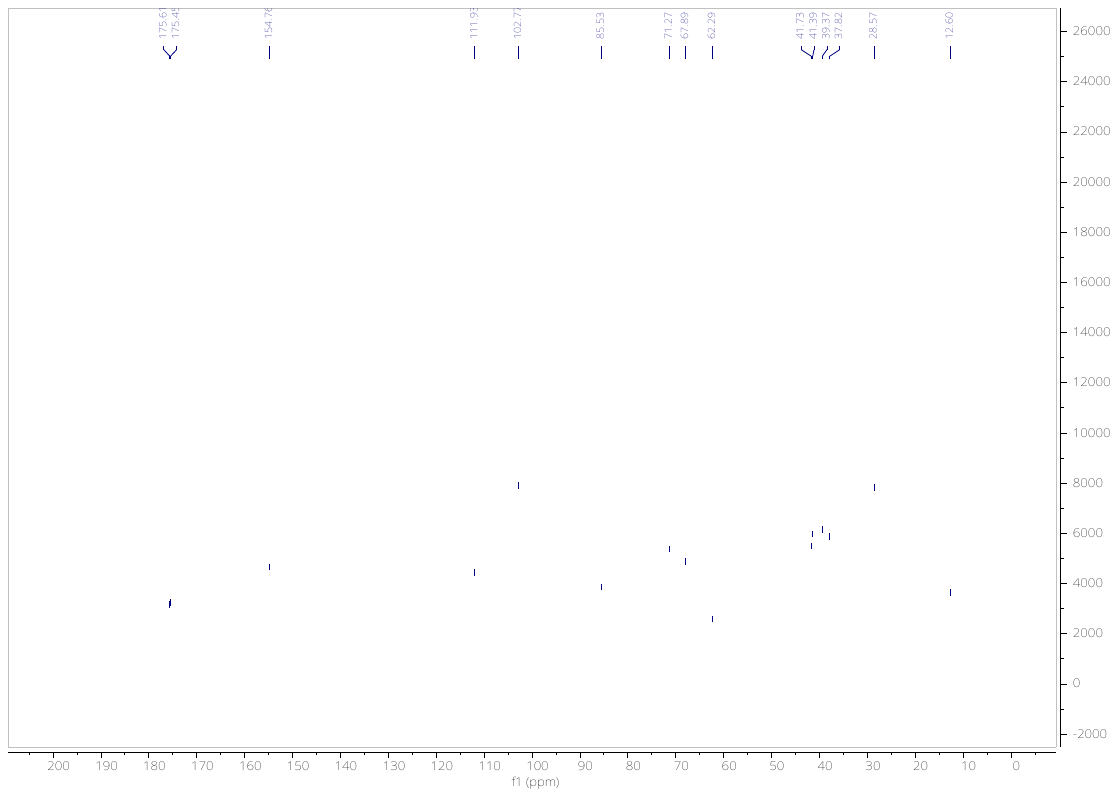

**Fig. S11.**^13^C NMR (400 MHz) Spectrum of Compound **3** in CD_3_OD

#### Figures S12-S15.1D and 2D NMR spectra of 4 in CD_3_OD

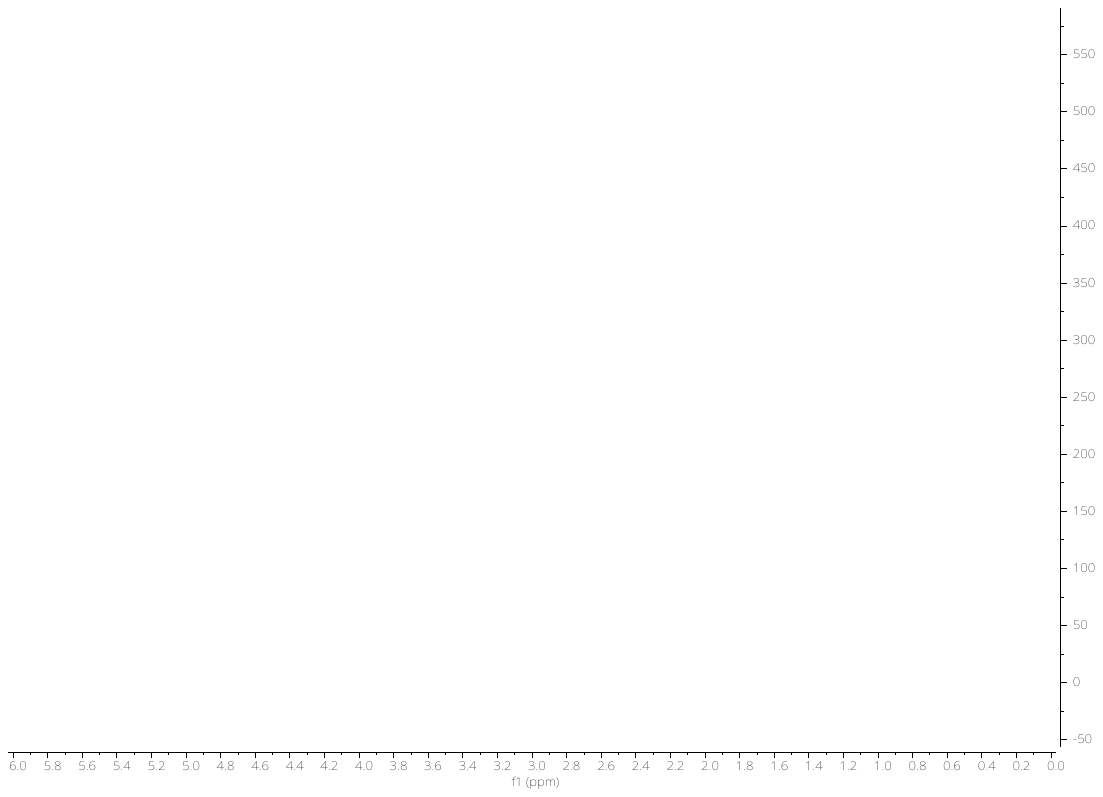

**Fig. S12.**^1^H NMR (400 MHz) Spectrum of Compound **4** (impurified with 35% of 3) in CD_3_OD

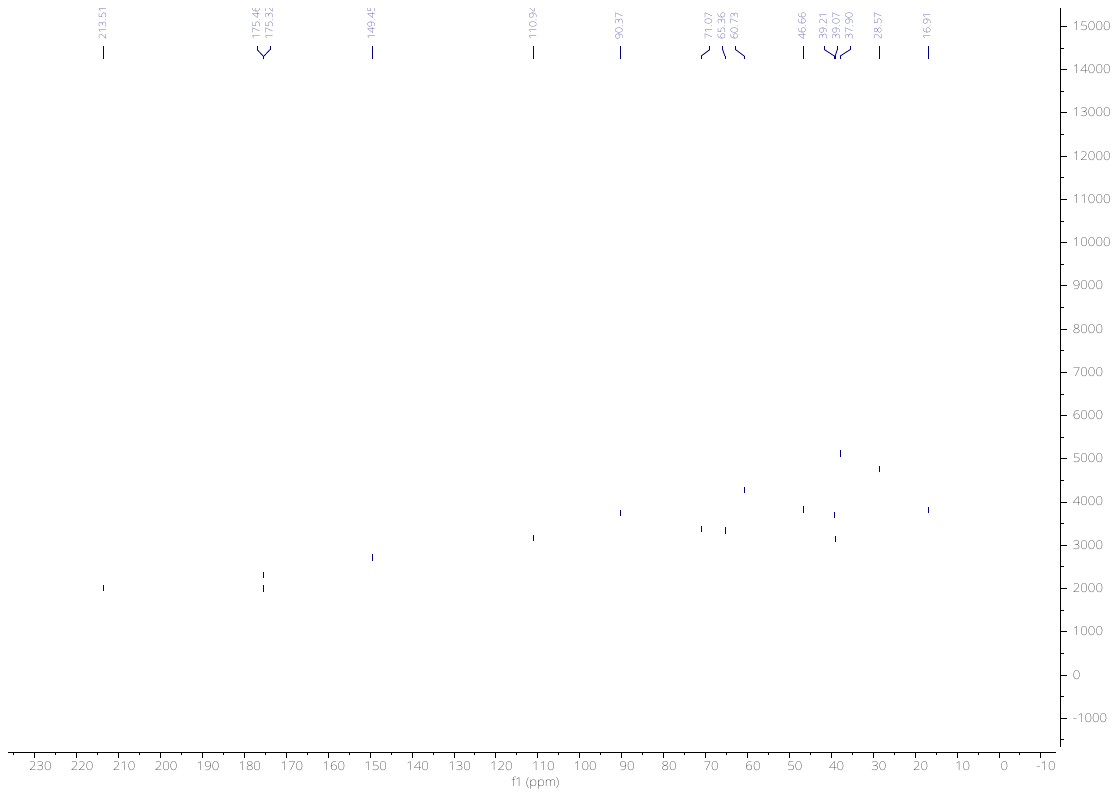

**Fig. S13.**^13^**C**NMR (400 MHz) Spectrum of Compound 4 (impurified with 35% of 3) in CD_3_OD

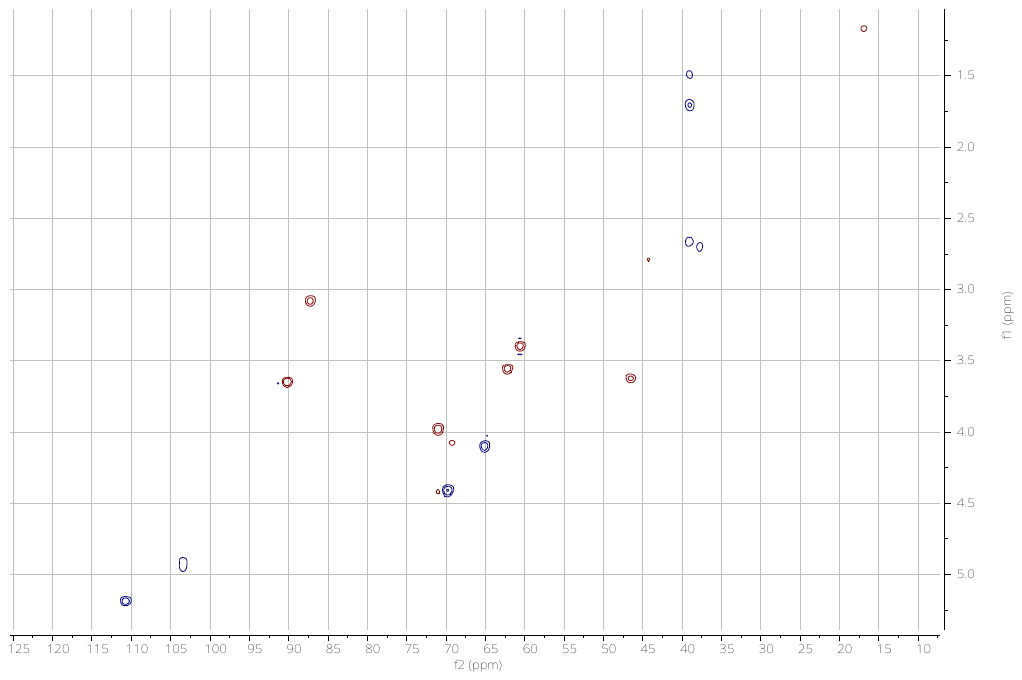

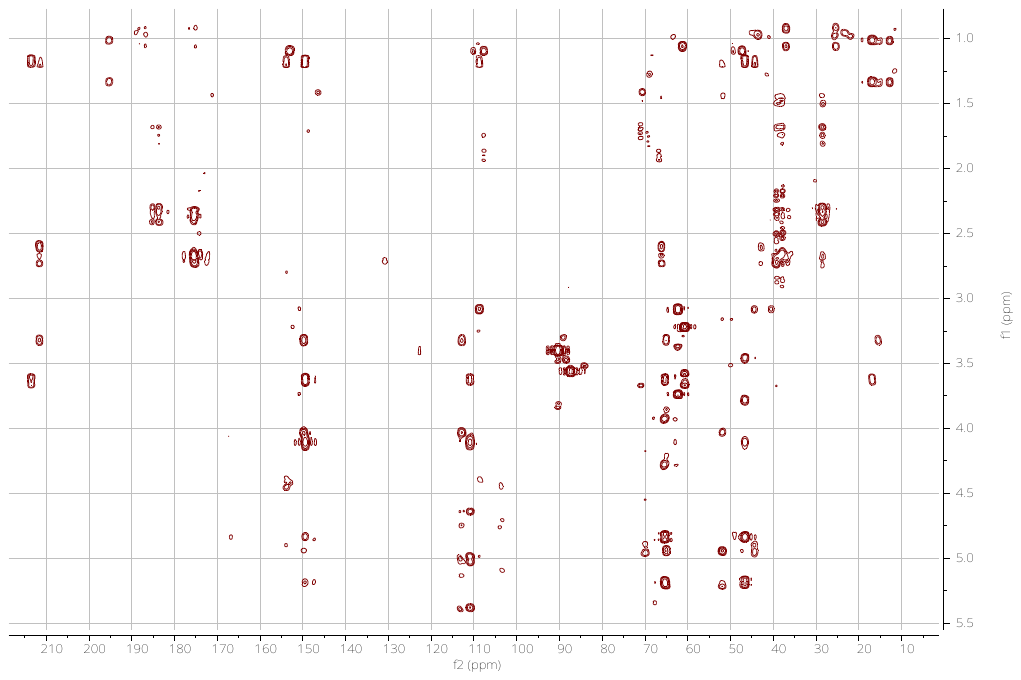

**Fig. S14.** gHSQC (400 MHz) Spectrum of Compound **4** (impurified with 35% of 3) in CD_3_OD

**Fig. S15.** gHMBC (400 MHz) Spectrum of Compound **4** (impurified with 35% of 3) in CD_3_OD

#### Figures S16-S21.1D and 2D NMR spectra of 5 in CDCl_3_

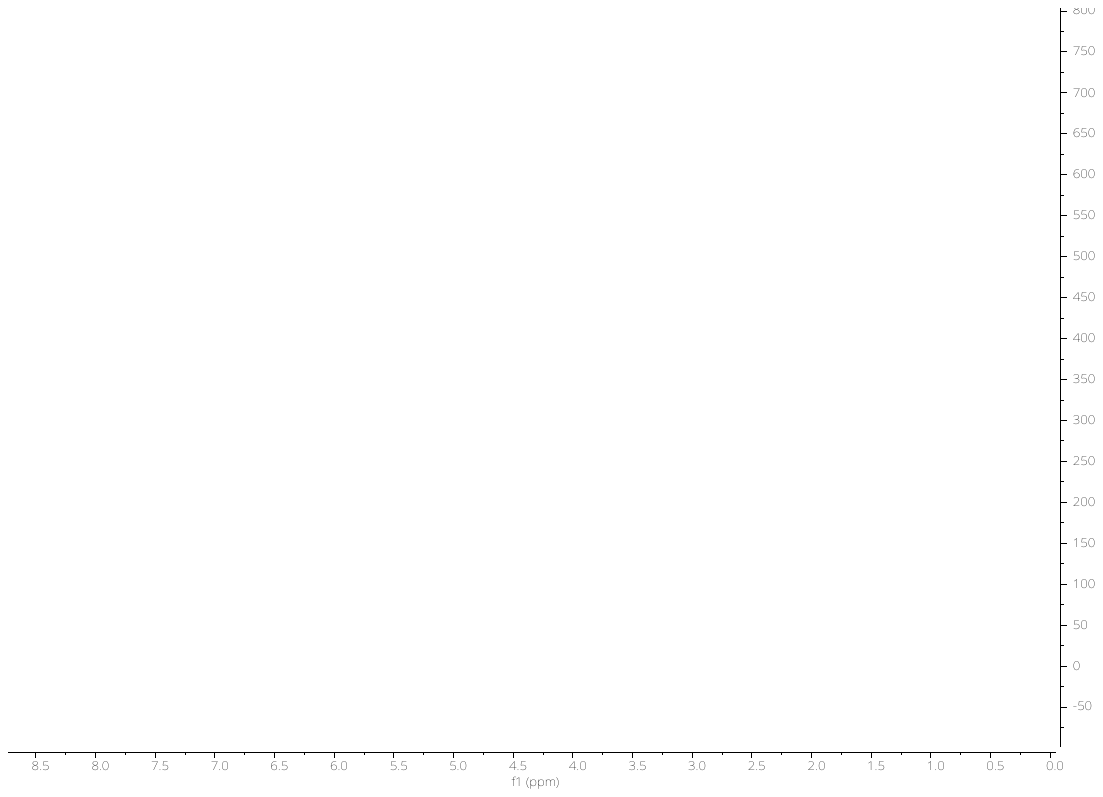

**Fig. S16.**^1^H NMR (500 MHz) Spectrum of Compound **5** in CDCl_3_

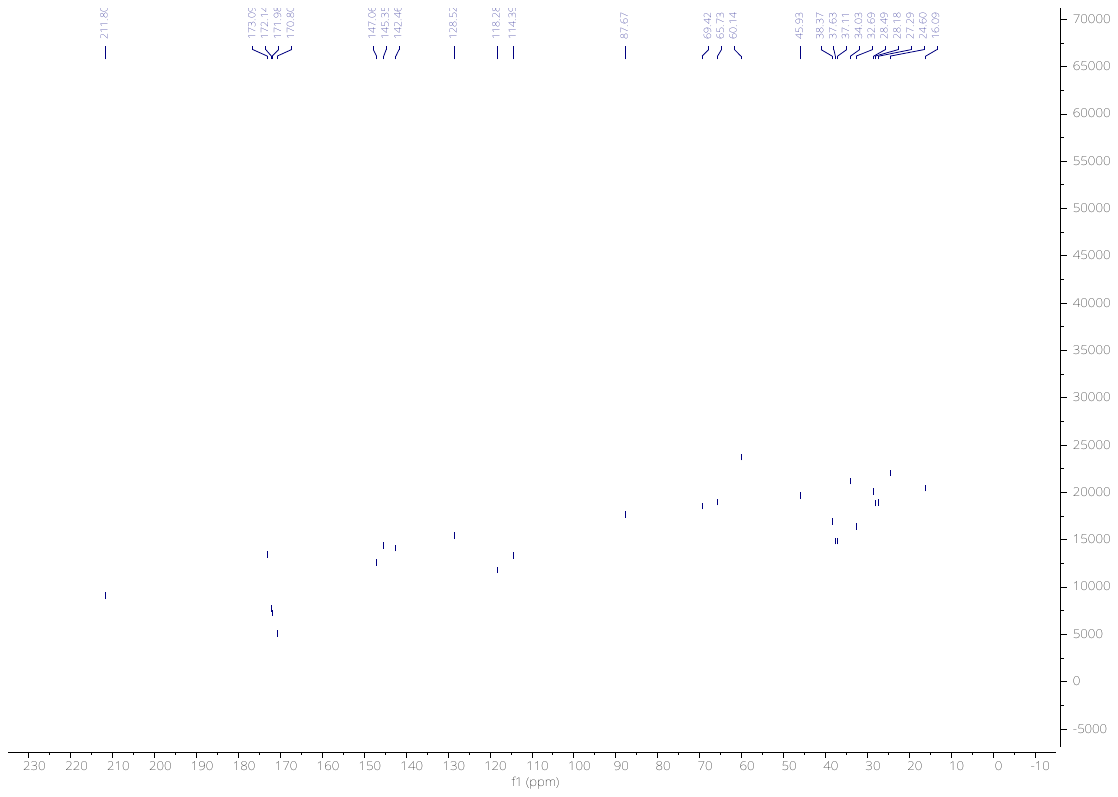

**Fig. S17.**^13^C NMR (100 MHz) Spectrum of Compound **5** in CDCl_3_

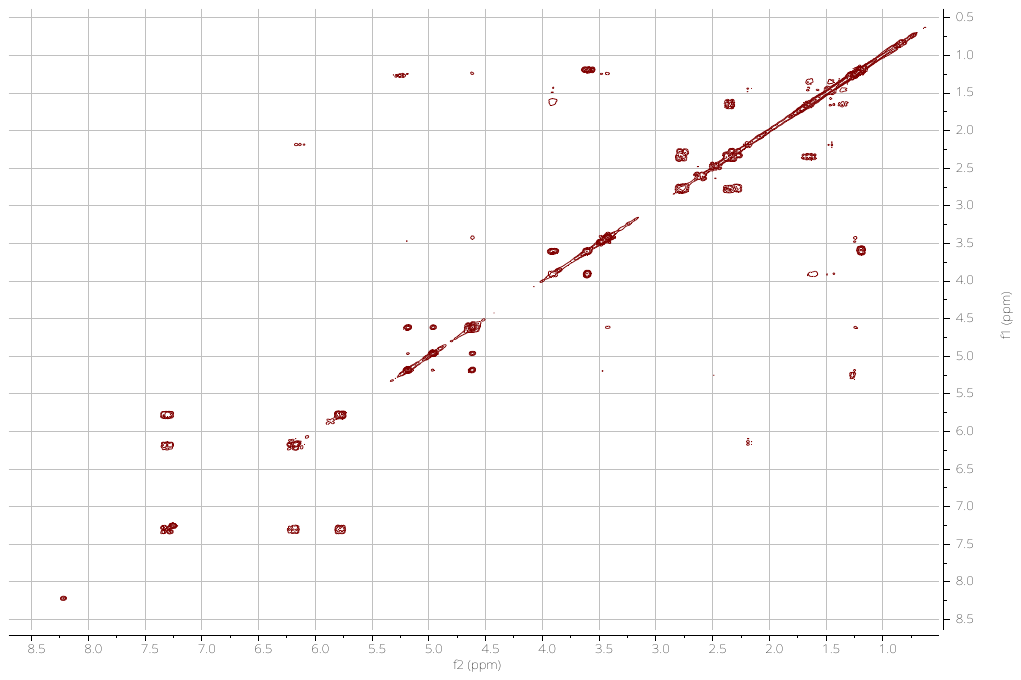

**Fig. S18.** gCOSY (500 MHz) Spectrum of Compound **5** in CDCl_3_

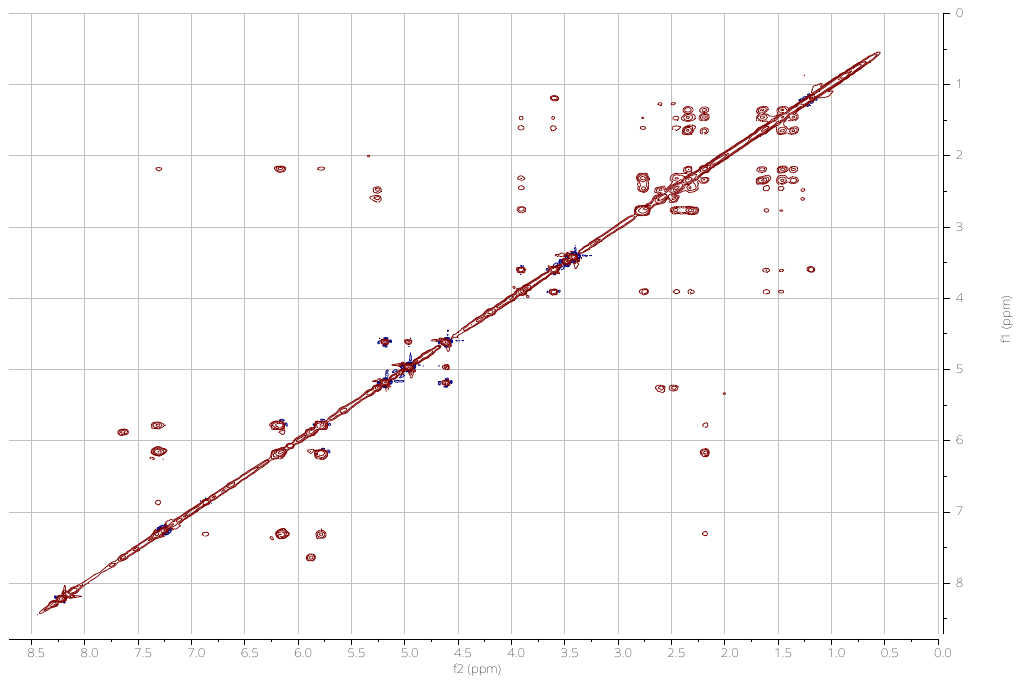

**Fig. S19.** TOCSY (500 MHz) Spectrum of Compound **5** in CDCl_3_

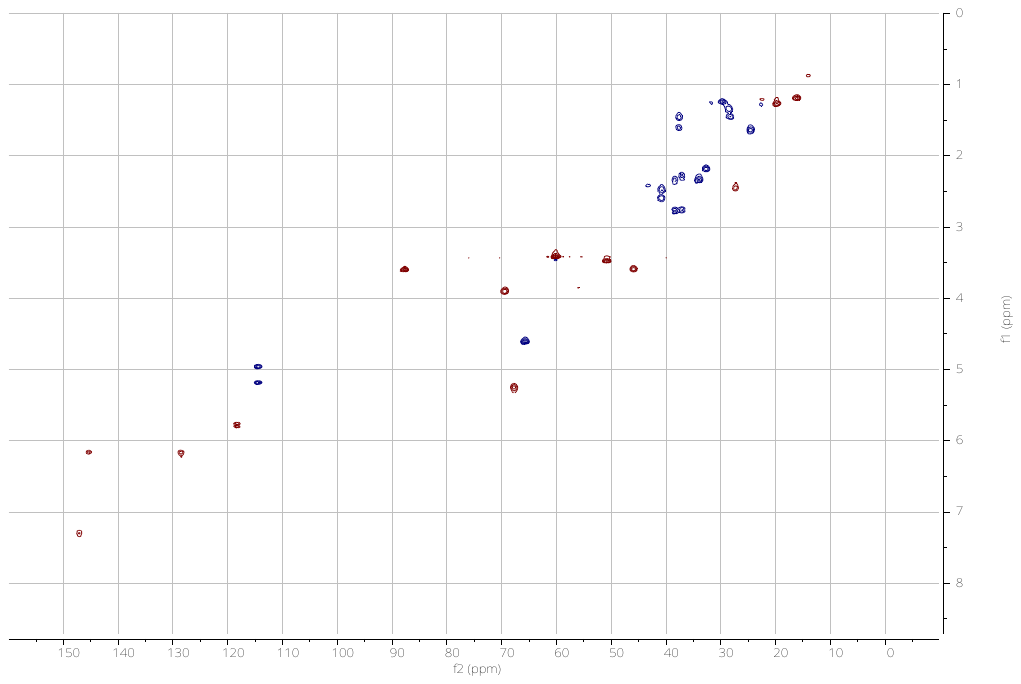

**Fig. S20.** gHSQC (500 MHz) Spectrum of Compound **5** in CDCl_3_

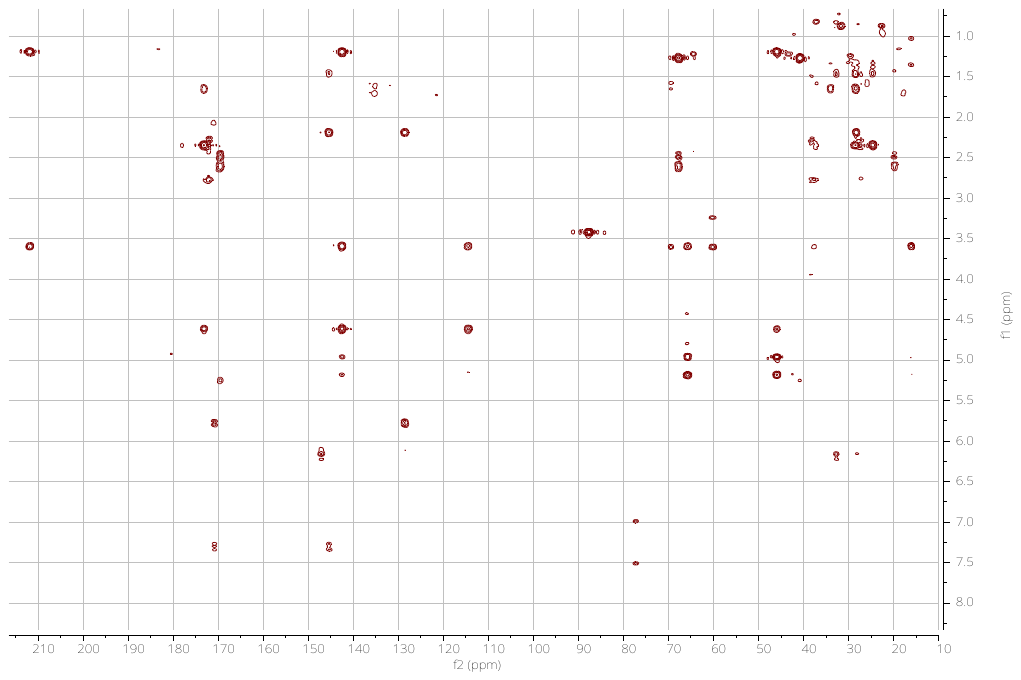

**Fig. S21.** gHMBC (500 MHz) Spectrum of Compound **5** in CDCl_3_

#### Figures S22-S24.ESIMS of 2, 4, 5

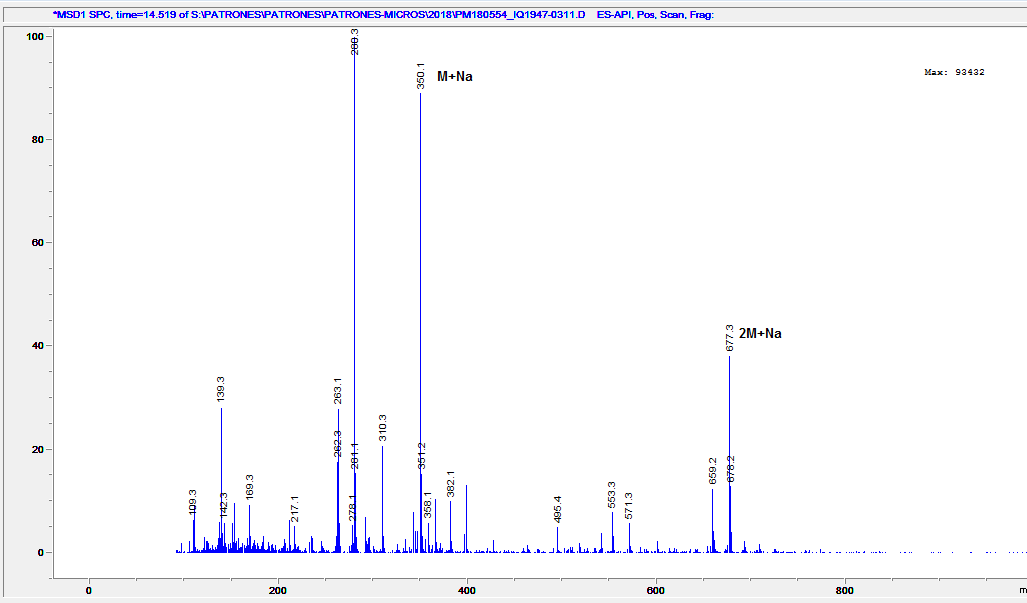

**Fig. S22.** ESIMS of compound **2**.

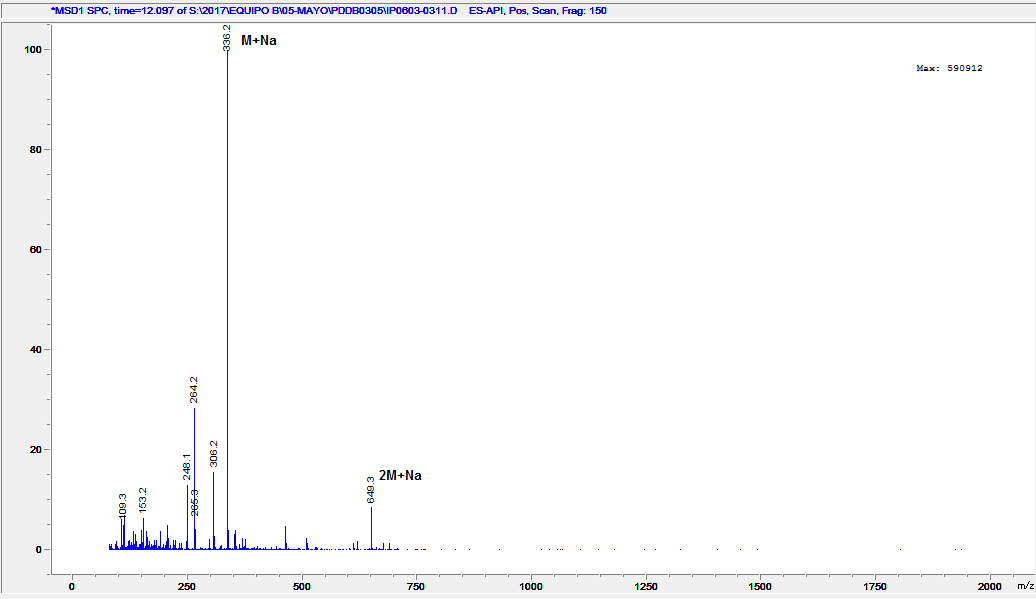

**Fig. S23.** ESIMS of compound **4**.

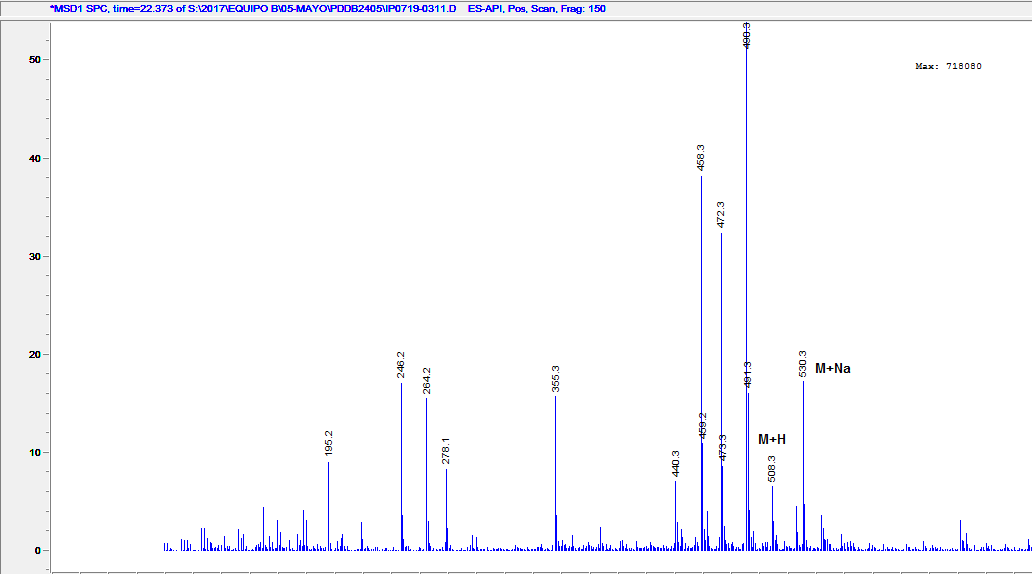

**Fig. S24.** ESIMS of compound **5**.

### Supplementary figures S25-S28

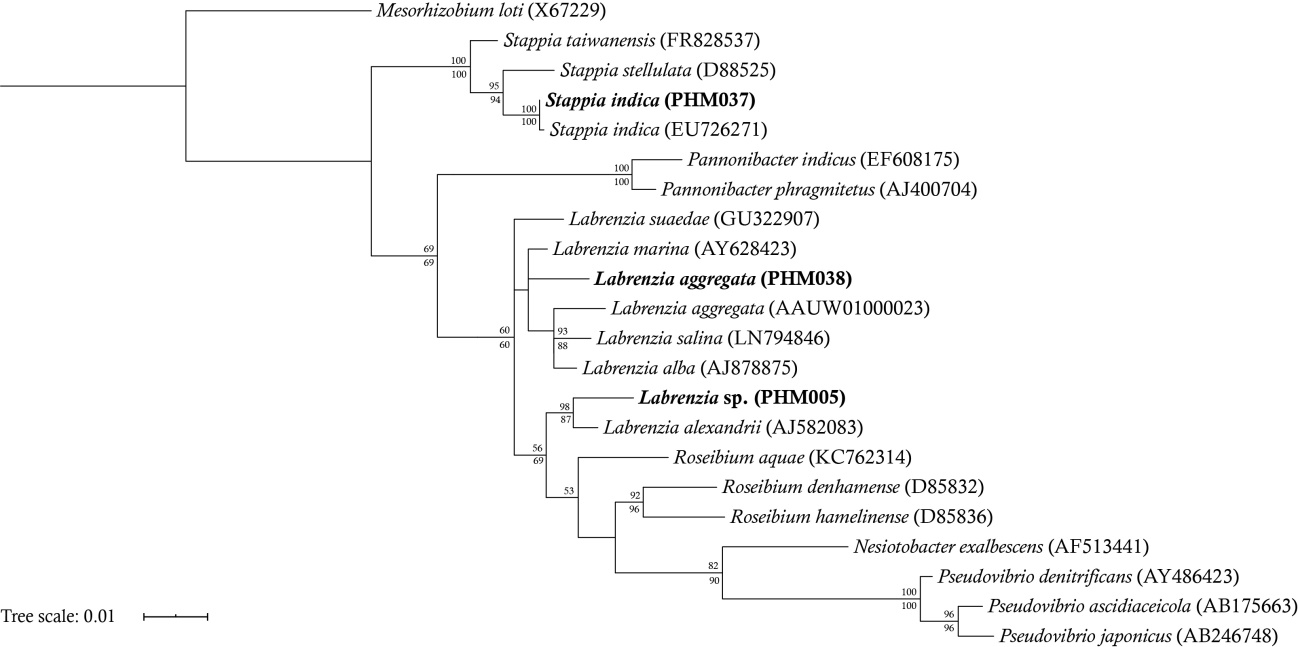

**Fig. S25.** Neighbor-joining (NJ) and Maximum Likelihood (ML) tree of 16S rRNA gene sequences reconstructed using TIM distances. In bold are included the samples sequenced for this work. The SILVA Accession numbers are indicated in parenthesis. Bootstrap values are from ML (above node) and NJ (below node) analyses. The tree is rooted with *Mesorhizobium loti*. SILVA accession numbers accompany each taxon name. The scale bar indicates evolutionary distances.

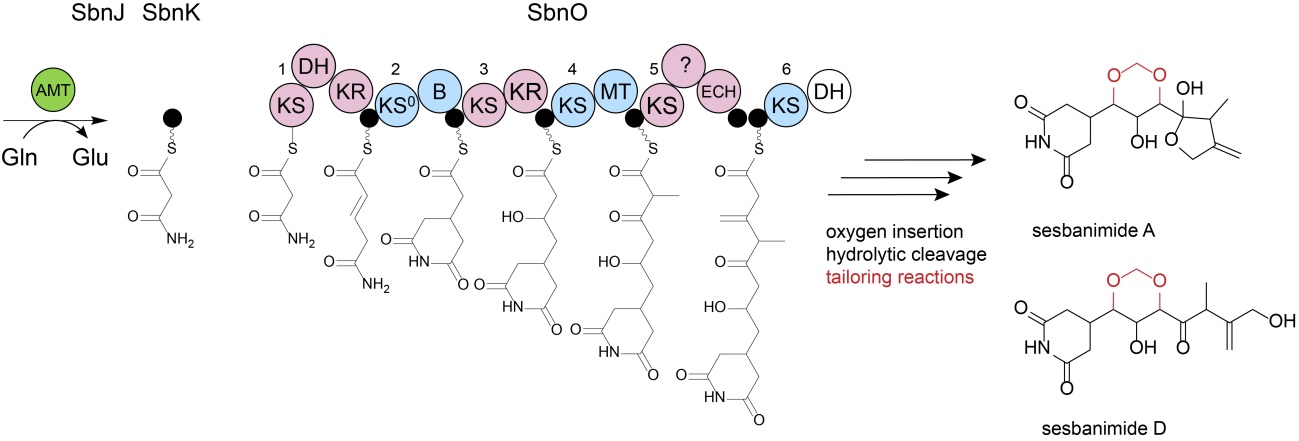

**Fig. S26.** Assembly of sesbanimide A analogs by PKS SbnO in *L. aggregata* PHM038. The loading module comprising amidotranferase (AMT) and associated acyl-carrier protein (ACP); SbnO. PKS domains indicated are ketoreductase (KR), dehydratase (DH), branching domain (B) and methyltransferase (MT). Different modules are numbered and indicated by the same color. White domains indicate supposed loss of activity. Tailoring reactions are marked in red.

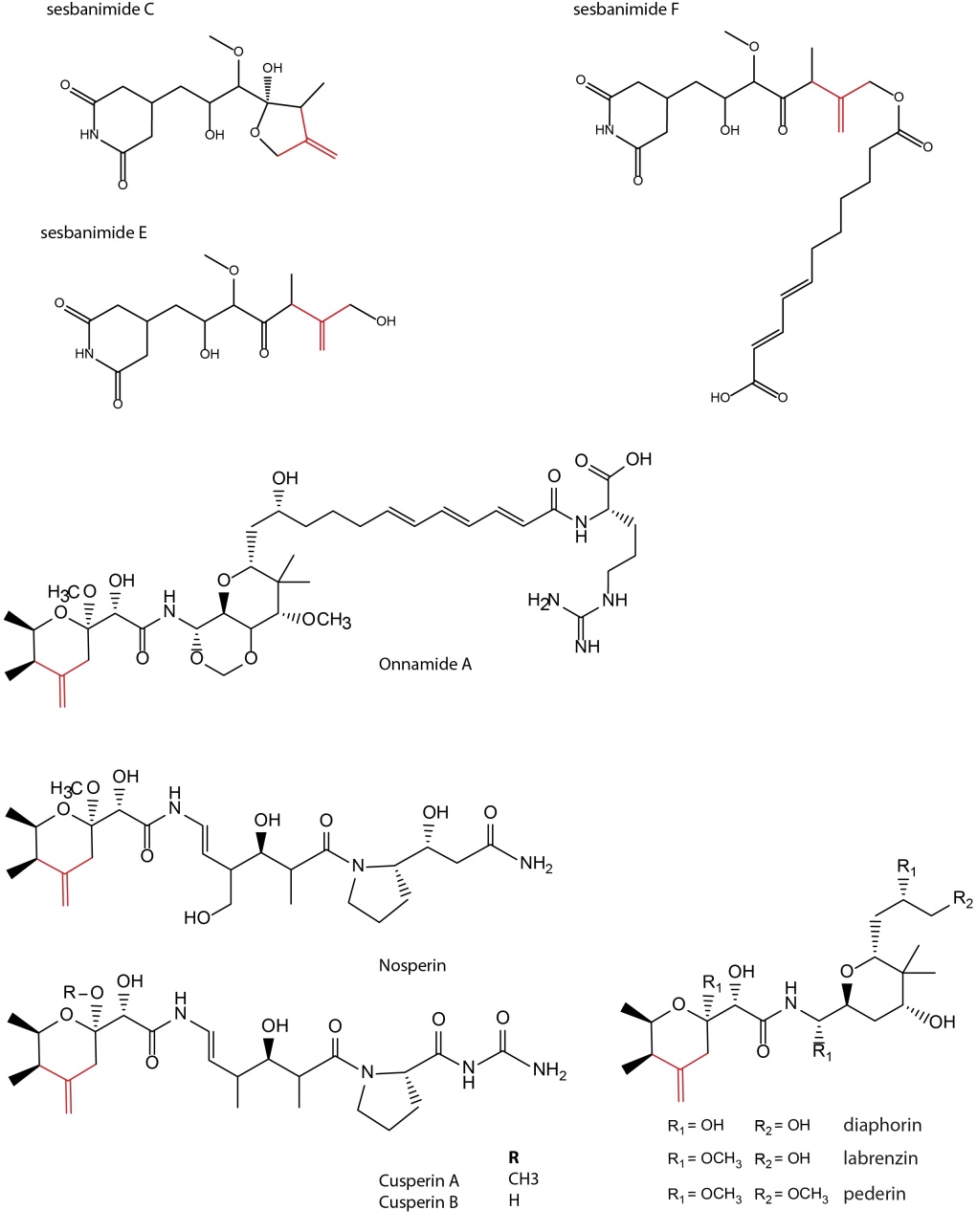

**Fig. S27.**Sesbanimide analogs, onnamide and pederin-like family compounds. The β-branching exomethylene group is colored in red.

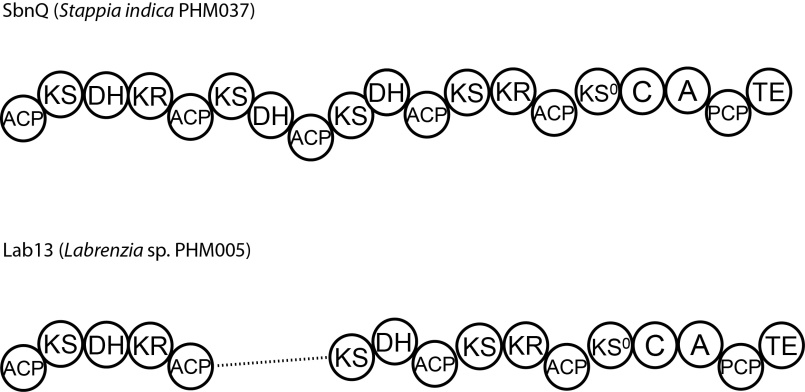

**Fig. S28.** Scheme to compare domain organization in Lab13 and SbnQ (PKS/NRPS).

### Supplementary tables

**Table S1.** Cluster for secondary metabolites in PHM037 and PHM038 detected by antiSMASH mining.

| strain | locus (bp) | clustertype | MIBiG homologue | Accession |
| --- | --- | --- | --- | --- |
| PHM037 | 5,035,472 - 5,076,482 | siderophore | deferrioxamine E | BGC0000940 |
|  | 4,939,412 - 5,017,119 | NRPS and type I PKS | micacocidin | BGC0001014 |
|  | 3,707,745 - 3,755,342 | NRPS | quinolobactin | BGC0000925 |
|  | 2,044,736 - 2,086,259 | phosphonate | phosphinothricintripeptide | BGC0000406 |
| PHM038 | NODE 31 (3,259 - 25,757) | betalactone | fengycin | BGC0001095 |
|  | NODE 22 (47,120 - 94,712) | bacteriocin |  |  |
|  | NODE 17 (126,880 - 147,339) | homoserinelactone |  |  |
| PHM037 and PHM038 | 4,845,006 - 4,891,839, PHM037; NODE 10 (47,120 - 94,712), PHM038 | type I PKS |  |  |
|  | 384,608 - 405,486, PHM037; NODE 8 (86,074 - 106,931), PHM038 | terpene |  |  |
|  | 1,731,410 - 1,746,233, PHM037; NODE 5 (92,449 - 107,188), PHM038 | N-acetylglutaminylglutamine amide |  |  |

**Table S2.** Known bacterial biosynthetic gene clusters containing genes homologous to those found in the *sbn* cluster.

| Characterizedclusters and hosts | NCBI accession | homologues in *sbn*cluster |
| --- | --- | --- |
| labrenzin cluster from *Labrenzia* sp. PHM005 | CP041191 | HMGS,ACP,KS,ECH,AT1,AT2,OXY, P450,PPT,MT,ABC |
| calyculin A cluster from uncultured Candidatus *Entotheonella* sp. | AB933566.1 | HMGS,ACP,KS,ECH,AT2 |
| macrobrevin cluster from *Brevibacillus* sp. Leaf182 | LMPN02000017.1 | HMGS,ACP,KS,ECH,AT2 |
| kalimantacin Aclusterfrom *Pseudomonas fluorescens* | GU479979.1 | HMGS,ACP,KS,ECH,AT2 |
| oocydin A cluster from *Serratia marcescens* | JX315603.1 | HMGS,ACP,KS,ECH,AT2,OXY |

**Table S3.** ORFs from the *sbn* cluster and their closest homologues in the RefSeq database.

| ORF | **Protein/**  **gene** | **putativefunction/homologue** | **Origin** | **Cover/identity %** | **Accession number of a protein homologue** |
| --- | --- | --- | --- | --- | --- |
| *SbnA* | **AT/AH** | Acyltransferase domain-containing protein | *Stappia indica* USBA 352 | 100/97.82 | [WP_067221761.1](https://www.ncbi.nlm.nih.gov/protein/WP_067221761.1?report=genbank&log$=prottop&blast_rank=2&RID=7M7G0NPD014) |
| *SbnB* | **4’-PPT** | 4'-phosphopantetheinyl transferase | *Stappia indica* USBA 352 | 99/98.17 | [WP_067221763.1](https://www.ncbi.nlm.nih.gov/protein/WP_067221763.1?report=genbank&log$=prottop&blast_rank=2&RID=7M7C8MZ3016) |
| *SbnC* | **EST** | [metallophosphoesterase](https://blast.ncbi.nlm.nih.gov/Blast.cgi#alnHdr_WP_120269009) | *Stappia indica* USBA 352 | 100/97.22 | [[WP_067221764.1](https://www.ncbi.nlm.nih.gov/protein/WP_067221764.1?report=genbank&log$=prottop&blast_rank=3&RID=7M7514PZ016)](https://www.ncbi.nlm.nih.gov/protein/WP_120269009.1?report=genbank&log$=prottop&blast_rank=2&RID=7M6FX11X014) |
| *SbnD* | **MT** | [FkbM family methyltransferase](https://blast.ncbi.nlm.nih.gov/Blast.cgi#alnHdr_WP_083202408) | *Stappia indica* USBA 352 | 100/94.03 | [WP_083202408.1](https://www.ncbi.nlm.nih.gov/protein/WP_083202408.1?report=genbank&log$=prottop&blast_rank=3&RID=7M6V1C0D014) |
| *SbnE* | **P450** | [cytochrome P450](https://blast.ncbi.nlm.nih.gov/Blast.cgi#alnHdr_WP_083206337) | *Stappia indica* USBA 352 | 100/97.61 | [WP_083206337.1](https://www.ncbi.nlm.nih.gov/protein/WP_083206337.1?report=genbank&log$=prottop&blast_rank=2&RID=7M7M2JNE014) |
| *SbnF* | **HMGS** | [hydroxymethylglutaryl-CoA synthas](https://blast.ncbi.nlm.nih.gov/Blast.cgi#alnHdr_WP_120269012)e | *Stappia* sp. ARW1T | 100/98.09 | [WP_120269012.1](https://www.ncbi.nlm.nih.gov/protein/WP_120269012.1?report=genbank&log$=prottop&blast_rank=2&RID=7M7SUE95014) |
| *SbnG* | **ACP** | Acyl carrier protein | *Stappia indica* USBA 352 | 100/98.77 | [WP_067221771.1](https://www.ncbi.nlm.nih.gov/protein/WP_067221771.1?report=genbank&log$=protalign&blast_rank=3&RID=7M7ZPFT0014) |
| *SbnH* | **KS** | [polyketide beta-ketoacyl:ACP synth](https://blast.ncbi.nlm.nih.gov/Blast.cgi#alnHdr_WP_120269013)ase | *Stappia* sp. ARW1T | 100/91.85 | [WP_120269013.1](https://www.ncbi.nlm.nih.gov/protein/WP_120269013.1?report=genbank&log$=prottop&blast_rank=2&RID=7M84XE9J014) |
| *SbnI* | **ECH** | [enoyl-CoA hydratase/isomerase](https://blast.ncbi.nlm.nih.gov/Blast.cgi#alnHdr_WP_158195488) | *Stappia indica* USBA 352 | 100/94.94 | [WP_067339094.1](https://www.ncbi.nlm.nih.gov/protein/WP_067339094.1?report=genbank&log$=prottop&blast_rank=2&RID=7M8AVU4R014) |
| *SbnJ* | **AMT** | [asparagine synthase (glutamine-hydrolyzing)](https://blast.ncbi.nlm.nih.gov/Blast.cgi#alnHdr_WP_067221776) | *Stappia indica* USBA 352 | 100/93.87 | [WP_067221776.1](https://www.ncbi.nlm.nih.gov/protein/WP_067221776.1?report=genbank&log$=prottop&blast_rank=3&RID=7M8E5G47016) |
| *SbnK* | **ACP** | [acyl carrier protein](https://blast.ncbi.nlm.nih.gov/Blast.cgi#alnHdr_WP_158195490) | *Stappia* sp. ARW1T | 100/96.34 | [WP_120269016.1](https://www.ncbi.nlm.nih.gov/protein/WP_120269016.1?report=genbank&log$=prottop&blast_rank=2&RID=7M8PGE5G014) |
| *SbnL* | **ABC** | cyclic peptide export ABC transporter | *Stappia indica* USBA 352 | 100/96.33 | WP_083202407.1 |
| *SbnM* | **ABC** | cyclic peptide export ABC transporter | *Stappia indica* USBA 352 | 100/97.27 | [WP_097175571.1](https://www.ncbi.nlm.nih.gov/protein/WP_097175571.1?report=genbank&log$=prottop&blast_rank=2&RID=7M96R96M014) |
| *SbnN* | **AT** | ACP S-malonyltransferase | *Stappia indica* USBA 352 | 100/94.66 | WP_097175572.1 |
| *SbnO* | **PKS** | [SDR family NAD(P)-dependent oxidoreductase](https://blast.ncbi.nlm.nih.gov/Blast.cgi#alnHdr_WP_120269020) | *Stappia* sp. ARW1T | 100/88.66 | WP_120269020.1 |
| *SbnP* | **OXY** | NAD(P)-binding domain-containing protein; monooxygenase | *Stappia indica* USBA 352 | 100/96.50 | [WP_097175574.1](https://www.ncbi.nlm.nih.gov/protein/WP_097175574.1?report=genbank&log$=prottop&blast_rank=2&RID=7M9UU1YB014) |
| *SbnQ* | **PKS/NRPS** | [non-ribosomal peptide synthetase](https://blast.ncbi.nlm.nih.gov/Blast.cgi#alnHdr_WP_067221787) | *Stappia indica* USBA 352 | 100/86.90 | [WP_067221787.1](https://www.ncbi.nlm.nih.gov/protein/WP_067221787.1?report=genbank&log$=prottop&blast_rank=2&RID=7MA35BKZ014) |
| *SbnR* | **HYP1** | [prohibitin family protein](https://blast.ncbi.nlm.nih.gov/Blast.cgi#alnHdr_WP_158195497) | *Stappia indica* USBA 352 | 100/97.92 | [WP_067221789.1](https://www.ncbi.nlm.nih.gov/protein/WP_067221789.1?report=genbank&log$=prottop&blast_rank=2&RID=7MABWPBX014) |
| *SbnS* | **ABC** | [ABC transporter substrate-binding protein](https://blast.ncbi.nlm.nih.gov/Blast.cgi#alnHdr_WP_158195498) | *Stappia indica* USBA 352 | 100/92.36 | [WP_067221790.1](https://www.ncbi.nlm.nih.gov/protein/WP_067221790.1?report=genbank&log$=prottop&blast_rank=2&RID=7MAGVPDY016) |
| *SbnT* | **HYP2** | Hypothetical protein | *Stappia indica* USBA 352 | 97/94.37 | [WP_067221792.1](https://www.ncbi.nlm.nih.gov/protein/WP_067221792.1?report=genbank&log$=prottop&blast_rank=2&RID=7MAPNK28014) |
| *SbnU* | **HYP3** | DUF697 domain-containing protein | *Stappia* sp.ARW1T | 100/97.17 | [WP_120269026.1](https://www.ncbi.nlm.nih.gov/protein/WP_120269026.1?report=genbank&log$=prottop&blast_rank=2&RID=7MAVM0ZC016) |
| *SbnV* | **HYP4** | DUF697 domain-containing protein | *Stappia* sp. ARW1T | 99/86.78 | [WP_147421864.1](https://www.ncbi.nlm.nih.gov/protein/WP_147421864.1?report=genbank&log$=prottop&blast_rank=2&RID=7MB044GM014) |
| *SbnW* | **HYP5** | [ParA](https://blast.ncbi.nlm.nih.gov/Blast.cgi#alnHdr_SOC15903) family protein | *Stappia indica* USBA 352 | 100/99.08 | WP_097175579.1 |

**Table S4.** Ketosynthase domain specificities and clade scores in the SbnO protein (PHM037 strain)

KS1

Clade_95 various specificities (E-value: 1.3E-173 Score: 570.7)
Clade_109 completely reduced (E-value: 6.8E-171 Score: 561.5)
Clade_8 unusual starter (AMT/Succinate) (E-value: 8.9E-216 Score: 709.6)
Clade_136 β D-OH (E-value: 4.4E-176 Score: 578.6)
Clade_96 various specificities (mainly α-Me) (E-value: 6.6E-172 Score: 565.0)

KS2

Clade_64 non-elongating (double bonds (mostly z-configured)) (E-value: 1.7E-214 Score: 705.2)
Clade_101 double bonds (E-value: 6.2E-146 Score: 479.3)
Clade_5 amino acids (oxa/thia) (E-value: 1.0E-145 Score: 478.7)
Clade_82 double bonds (mostly e-configured) (E-value: 5.0E-145 Score: 476.4)
Clade_115 β-keto or double bonds (E-value: 1.2E-143 Score: 471.7)

KS3

Clade_95 various specificities (E-value: 5.5E-167 Score: 548.8)
Clade_96 various specificities (mainly α-Me) (E-value: 4.7E-188 Score: 618.2)
Clade_12 vinylogous chain branching (E-value: 3.1E-183 Score: 602.2)
Clade_136 β D-OH (E-value: 2.3E-165 Score: 543.3)
Clade_40 β D-OH or β-keto (E-value: 1.8E-159 Score: 523.9)

KS4

Clade_7 ß D-OH (E-value: 2.4E-215 Score: 708.4)
Clade_110 ß D-OH or double bonds (e-configured) (E-value: 2.3E-221 Score: 728.0)
Clade_140 β D-OH (E-value: 4.3E-214 Score: 704.3)
Clade_137 β D-OH (E-value: 5.9E-214 Score: 703.7)
Clade_62 β D-OH (some with α L-Me) (E-value: 1.9E-208 Score: 685.4)

KS5

Clade_104 ß O-Me or ß-Me double bond (E-value: 2.4E-165 Score: 543.0)
Clade_68 α L-OH/Me β D-OH (E-value: 4.6E-164 Score: 539.1)
Clade_21 α-Me reduced/keto/D-OH (E-value: 4.1E-181 Score: 595.2)
Clade_74 α-Me reduced/keto/D-OH (E-value: 1.2E-178 Score: 587.1)
Clade_23 α-Me (E-value: 3.7E-175 Score: 575.5)

KS6

Clade_104 ß O-Me or ß-Me double bond (E-value: 7.2E-165 Score: 541.5)
Clade_86 α-L-Me red or OH (E-value: 1.1E-159 Score: 524.6)
Clade_14 exomethyl/exoester (E-value: 5.5E-189 Score: 621.2)
Clade_2 α-Me shifted double bond or OH (E-value: 2.3E-168 Score: 553.1)
Clade_73 exomethylene (E-value: 1.7E-157 Score: 517.2)

**Table S5**. Ketosynthase domain specificities and clade scores in the SbnQ protein (PHM037 strain)

KS1

Clade_35 oxidative rearrangement (E-value: 2.4E-220 Score: 724.8)
Clade_95 various specificities (E-value: 2.9E-173 Score: 569.5)
Clade_25 completely reduced (E-value: 4.9E-178 Score: 585.0)
Clade_136 β D-OH (E-value: 6.9E-177 Score: 581.2)
Clade_96 various specificities (mainly α-Me) (E-value: 4.6E-176 Score: 578.7)

KS2

Clade_95 various specificities (E-value: 3.3E-183 Score: 602.2)
Clade_25 completely reduced (E-value: 1.2E-211 Score: 695.8)
Clade_108 shifted double bonds (E-value: 3.9E-199 Score: 654.7)
Clade_96 various specificities (mainly α-Me) (E-value: 7.8E-189 Score: 620.8)
Clade_11 shifted double bonds (E-value: 4.8E-185 Score: 608.3)

KS3

Clade_25 completely reduced (E-value: 9.1E-218 Score: 716.0)
Clade_108 shifted double bonds (E-value: 3.2E-206 Score: 678.0)
Clade_11 shifted double bonds (E-value: 3.7E-194 Score: 638.3)
Clade_96 various specificities (mainly α-Me) (E-value: 8.5E-188 Score: 617.3)
Clade_40 β D-OH or β-keto (E-value: 5.0E-187 Score: 614.7)

KS4

Clade_82 double bonds (mostly e-configured) (E-value: 2.7E-222 Score: 731.0)
Clade_125 double bonds (e-configured) (E-value: 8.8E-220 Score: 722.7)
Clade_115 β-keto or double bonds (E-value: 7.2E-214 Score: 703.1)
Clade_129 double bonds (e-configured) (E-value: 8.9E-210 Score: 689.8)
Clade_101 double bonds (E-value: 1.7E-209 Score: 688.7)

KS5

Clade_76 non-elongating (double bonds) (E-value: 6.7E-172 Score: 564.9)
Clade_142 non-elongating (various) (E-value: 1.1E-161 Score: 531.1)
Clade_101 double bonds (E-value: 3.0E-158 Score: 519.9)
Clade_90 β-keto or double bonds (E-value: 7.3E-158 Score: 518.8)
Clade_128 double bonds (e-configured) (E-value: 4.5E-155 Score: 509.5)
